# DemuxHMM: large-scale single-cell embryo profiling via recombination barcoding

**DOI:** 10.64898/2026.02.23.703392

**Authors:** Anton Afanassiev, Kevin Wei, Nozomu Yachie, Kenji Sugioka, Geoffrey Schiebinger

## Abstract

High-resolution developmental time-courses with single-cell RNA sequencing (scRNA-seq) increasingly target trajectory inference and other analyses in the study of development and disease. Researchers can efficiently generate these datasets by pooling individuals and inferring cell-to-individual mappings after sequencing, in a process called demultiplexing. Existing demultiplexing methods face limitations in the number of timepoints they can support, due to either the need for individual-by-individual processing or reduced accuracy at large numbers of individuals. To address these limitations, we introduce a computational framework for demultiplexing large-scale, individual-resolved datasets. Our framework includes a recombination-aware demultiplexing method, DemuxHMM, which uses a Hidden Markov Model (HMM) to explicitly model contiguous SNP patterns that arise from simple breeding schemes. We demonstrate substantial performance and scalability gains from this recombination-aware approach on both a real Diversity Outbred (DO) mouse data and a simulated Drosophila dataset. This performance demonstrates the value of the method when used for common applications on model organisms such as time series, crossover position inference, and eQTLS, among others.

**Author summary:** To study aspects of various species, researchers measure ribonucleic acid (RNA) expression at the single-cell level for increasingly large cohorts of individuals. When the number of individuals is very large, mapping cells back to the individual of origin (demultiplexing) using lab techniques is costly and time-consuming. An emerging class of methods addresses this by using natural variation between individuals as a type of barcode to demultiplex. We create an algorithm in this class which works in scenarios where researchers breed organisms in defined cross schemes, which are common among model organisms. The breeding approach creates predictable patterns in the barcodes that our algorithm can exploit for greater performance and scalability. In this paper, we show that the approach works well on a real mouse dataset and compare it against competing methods on a rigorous simulated data test suite.

## 1 Introduction

The past decade has witnessed an explosion of single-cell datasets. While older technologies required processing cells in plates [1], where researchers had to manually manipulate each individual cell, new single-cell genomics technologies can process cells in *parallel* [2], enabling high-throughput profiling of large populations of cells.

The natural next step is to profile cells from many *individuals*, and explore how cells change over time, with disease, or across the population (e.g. with genetic background). In developmental time-courses, the ‘size’ of a dataset depends on the number of timepoints (i.e. individuals) collected, whereas the number of cells sampled determines the noise level of each timepoint along a developmental curve [3, 4]. Population-level analyses also gain statistical power from more individuals. These analyses include inferring natural selection based on expression differences [5] or detecting associations between those gene expression differences and their genetic determinants through eQTLS [6]. Crossover position mapping also requires large numbers of individuals [7]. However, profiling cells from many individual organisms requires some strategy for parallel processing of individuals.

Current techniques for parallel processing of individuals rely on pooling cells from individuals, sequencing the pool, and then using demultiplexing techniques to re-assign cells to their individual of origin. These demultiplexing techniques use either molecular barcodes [8–14], natural variation combined with collected genotypes [15–17], or are natural-variation-based and self-genotyping [18–21] (see Appendix 5.1 for a survey). The first two categories require individual-level processing to add tags or collect genotypes, limiting scale due to the labor and monetary costs involved. Self-genotyping, natural-variation-based methods instead focus on inferring identities of individuals using lossy “barcodes” of single nucleotide polymorphisms (SNPs), requiring no individual-level pre-processing. As such, self-genotyping methods position themselves as the most scalable class of approaches. Currently, they comprise four main methods: Vireo [18], Souporcell3 [19], scSplit [20], and freemuxlet [21]. However, each of these existing approaches treats SNPs as independent units, ignoring the fact that inheritance and recombination create chromosome-level structure. This treatment limits their accuracy and scalability when researchers pool many individuals or few cells are available per individual.

In this paper, we develop a demultiplexing method for parallel processing of single-cell data from large cohorts of individuals. Our method, DemuxHMM, leverages linkage in SNPs with a Hidden Markov Model (HMM) to cluster cells into individuals. At the same time, we also genotype each individual in the pool. Our approach enables accurate demultiplexing at the scales required for massive time series and other applications, without any individual-level handling.

One of the main goals of the paper is to demonstrate that DemuxHMM is actually feasible on real data of the scale that would be useful to the applications we mentioned. Our primary example is a 77-individual Diversity Outbred (DO) mouse dataset from Dillard et al. [22], that, after pre-processing, has over 86,000 cells. The large number of individuals and cells makes this dataset challenging for existing demultiplexing methods. At the same time, the DO breeding line is relevant to many of the aforementioned applications [23–26]. Subsequently, we present more comprehensive benchmarks on simulated *Drosophila melanogaster* datasets. In these benchmarks, we explore the limits of scalability, robustness, as well as doublet detection performance. Overall, the results reinforce the feasibility of DemuxHMM for demultiplexing.

## 2 Results

Viewed in terms of SNPs, a genotype serves as an individual-level barcode. Demultiplexing methods that infer these genotypes without prior knowledge need no individual-level handling. However, existing methods view SNPs as independent of each other [15–21]. This approach can be a good simplification in unrelated individuals, but it discounts the fact that SNPs follow a structured arrangement when individuals are genetically related (as illustrated in Figure 1).

**Fig 1.**
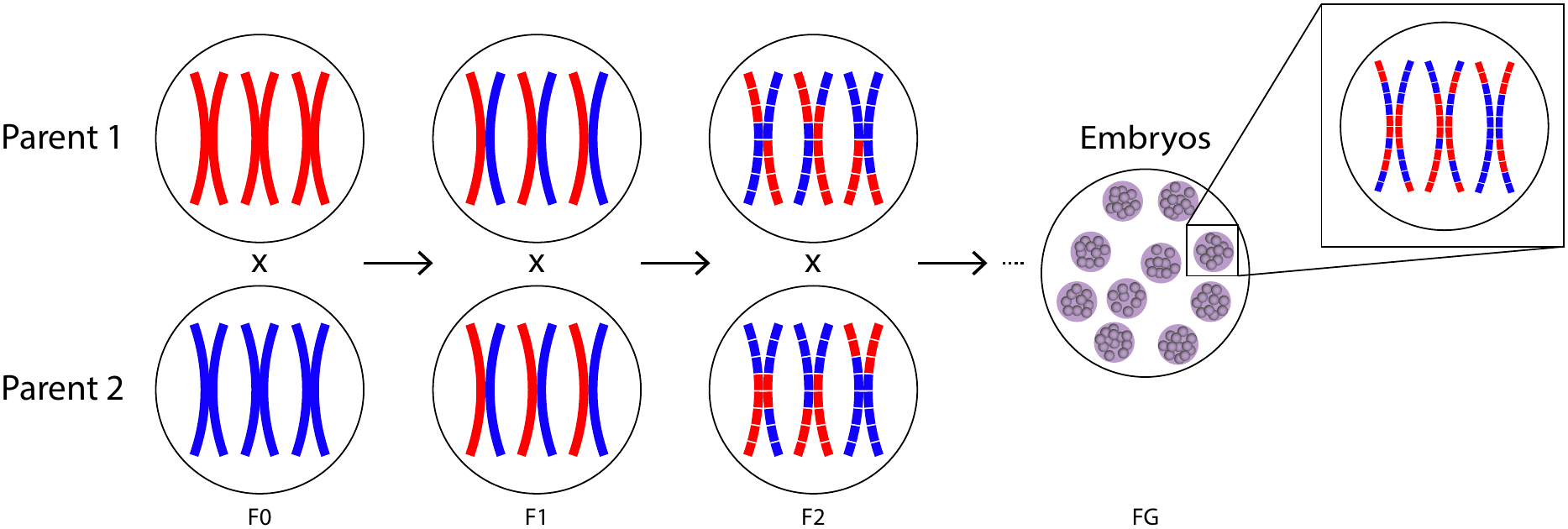
Single nucleotide polymorphism patterns provide noisy embryo barcodes. Consider two highly divergent parents (red and blue). These should vary at many polymorphic sites. In the figure, we visualize the parents as homozygous variant (red) and homozygous reference (blue), but note that homozygosity is not a requirement. The parents breed to create a pool of offspring, which interbreed to make further generations. At each generation, meiotic recombination through crossover events increases diversity, creating complex, but segmentally structured, patterns of variants in the offspring. This creates a pool of individuals that is highly diverse, but with an underlying structure that can be treated as a barcode.

Consider two parents from a model species, one treated as a reference strain and one from a highly divergent strain with respect to the first parent. The F1 generation inherits one chromosome from each parent. During the creation of gametes, meiotic recombination causes entire chromosomal segments to cross over. Thus, the F2 generation can have three possible genotypes at a chromosome location: homozygous reference, heterozygous, and homozygous variant (*g* ∈ {0, 1, 2}). These genotypes occur in contiguous segments of shared ancestry along each chromosome. Continuing breeding along further generations increases the complexity of this structure, creating informative, highly structured patterns of variation that existing demultiplexing methods do not exploit. Fig. 1 visualizes this breeding scheme in terms of parental genotypes. Furthermore, one can add time series structure in many ways, such as combining or mixing generations. In general, one can create contiguous SNPs segments with various other experimental structures. For example, Diversity Outbred mice use 8 diverse founding parents [27], and as we show in Sec. 2.2 this setup is also compatible with DemuxHMM. Overall, the contiguous SNP structure needed can be created in many ways.

DemuxHMM explicitly leverages contiguous SNP structure by modeling each chromosome as a Hidden Markov Model (HMM) on genotypes, with transitions governed by recombination-based probabilities. This approach allows DemuxHMM to capture recombination barcodes for individuals. Based on this strategy, the method infers the genotypes of individuals and then assigns cells to their most likely individual of origin based on observed variant profiles. By considering recombination barcoding in its demultiplexing framework, DemuxHMM performs strongly in challenging demultiplexing regimes, such as those with few cells per individual or large numbers of individuals. One can also use the method without any experimental prerequisites when transition probabilities are set to uniform, achieving comparable performance to existing self-genotyping methods. We demonstrate performance on an existing real mouse dataset with the related Diversity Outbreeding [27] structure in Sec. 2.2. We also provide more comprehensive benchmarks on a simulated *Drosophila melanogaster* model in Sec. 2.3.

### 2.1 A Recombination Driven Hidden Markov Model on Genotypes for Demultiplexing

In this section, we describe the mechanics of DemuxHMM, a demultiplexing approach that can leverage structured recombination. For input, the method expects users to have produced a pool of individuals (for example, a pool of embryos of different developmental timepoints) as described in Section 1 and Fig. 1. Users then sequence the pool, and apply subsequent quality control. They then perform variant calling, creating the primary input to the algorithm. Specifically, the data should be a collection of matrices *A*^*c*^, and *D*^*c*^, for each chromosome *c*. These matrices are of shape *N × M*_*c*_, where *N* is the number of collected cells, and *M*_*c*_ is the number of SNPs seen on chromosome *c*. 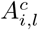 represents the number of variant counts seen for cell *i* at SNP position *l*, and 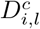 is the total number of counts seen at that position (depth). As output, the algorithm produces a clustering of cells to individuals along with inferred genotypes for each detected individual in the pool.

At its core, our approach fits a genotype to each individual and then assigns cells to it through maximum likelihood. Let there be *K* total individuals. Then, consider a chromosome *c* having *M*_*c*_ variants in individual *k*. We represent the chromosome with a chain of genotypes, 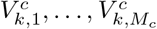 representing, in order, each detected SNP position on the chromosome. We assign each position a state in 0, 1, 2. The states represented are homozygous reference, heterozygous, and homozygous variant, respectively. We assume that the probability of transitioning between states is approximately independent of inter-SNP distance. In testing, we have found that incorporating inter-SNP distances does not result in large improvements in accuracy, suggesting that this is a reasonable assumption. The transition matrix *T*^*c*^ for each chromosome *c* governs the likelihood of transitioning between states. To fit the genotype model to the data, we use a similar approach to that of Vireo [18]. That is, we model the observed allele counts with a binomial distribution based on the state and its expected allele frequency. We update the variables of the model iteratively through a series of Expectation Maximization (EM) iterations until the model converges or we reach the maximum number of iterations. To determine convergence, we compute a truncated objective function at each iteration (defined in Section 4.2). We say the model has converged when the change in the objective shrinks below a user-input threshold. To summarize, Fig. 2A presents a high-level overview of the approach.

**Fig 2.**
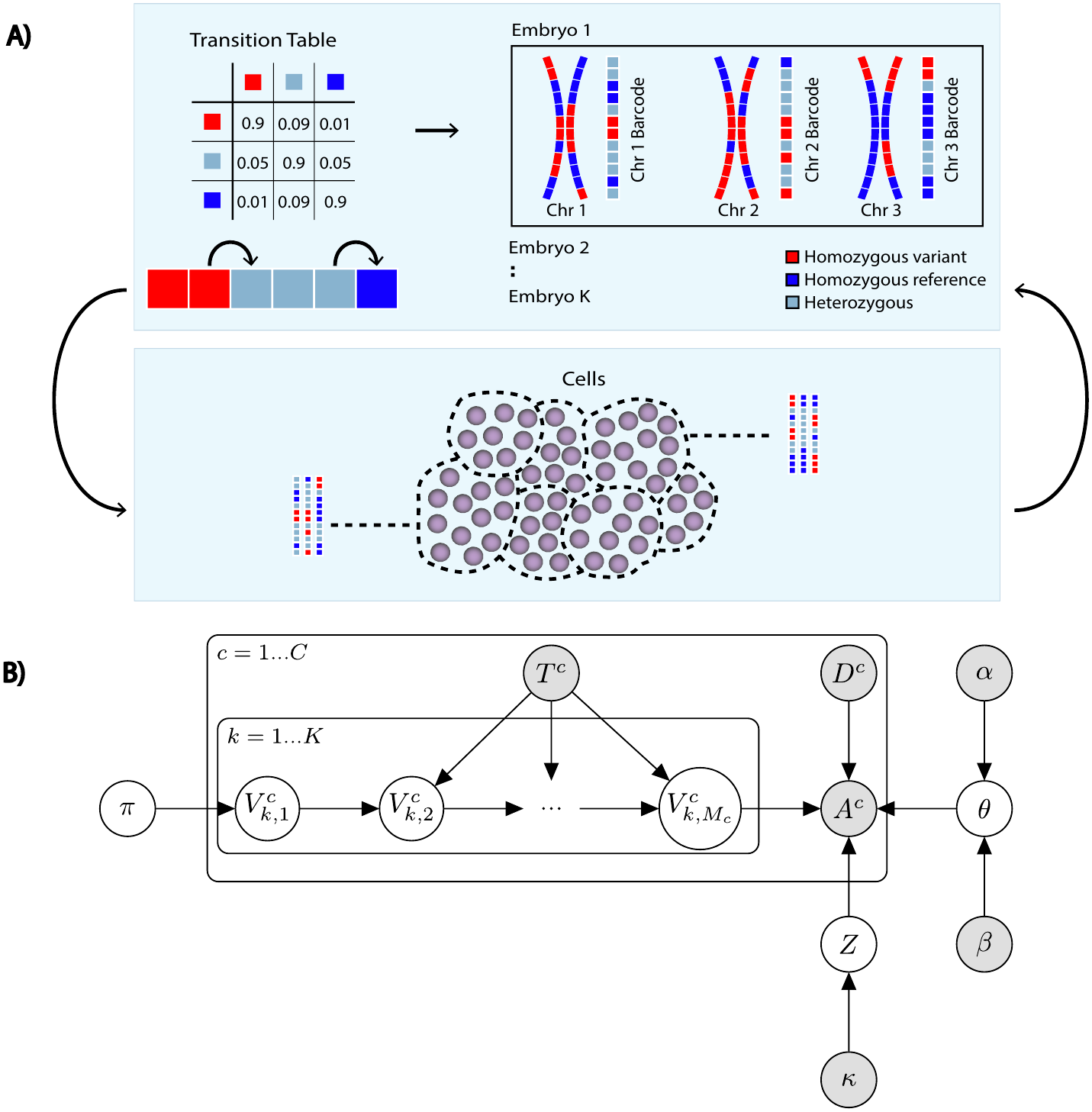
An overview of the DemuxHMM algorithm and model. In **A)** the demultiplexing stage of the DemuxHMM algorithm is shown. The algorithm takes variant calling data from a pool of individuals as input and produces a demultiplexing through EM iterations. Most importantly, DemuxHMM infers the unique pattern of variants that act as the “recombination barcode” for each chromosome of each individual. DemuxHMM uses a transition matrix to capture segmentally defined variant structure. This allows for greater performance when the method is combined with the suggested mating approach. The inference alternates between genotyping, and optimization of other variables, namely cell assignments. After each set of iterations, cell assignments and genotypes improve. **B)** A graphical representation of the DemuxHMM model. Shaded variables are input parameters. We infer unshaded variables during the optimization (latent). Arrows show dependency relationships. At the core of the model are the genotype encoding chains 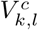. There are *M*_*c*_ (number of detected variants) of these for each chromosome, for each individual. *π* is a vector of length 3 holding initialization probabilities for 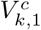. *T*^*c*^ is a 3 *×* 3 matrix holding transition probabilities for the chains at chromosome *c. A*^*c*^ is a matrix of size *N × M*_*c*_, where there are *N* collected cells. 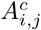 contains the number of counts seeing the alternative allele for variant *j*, for cell *i*, on chromosome *c*. 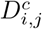 contains the total number of counts at that position (depth). *Z* is a vector of length *N* containing cell assignments, i.e. the result of the demultiplexing. It has prior *κ*, which is a probability vector of length *K*, where *K* is the expected number of individuals. Finally, *θ* is a vector of length 3, where *θ*_*t*_ represents the variant allele frequency of genotype *t*. It has a Beta prior with parameters *α, β*.

We now briefly look at the mathematical details of the model. First, we examine the chains of genotypes which encode the “barcodes” for each chromosome of each individual. In each chain we initialize 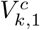 by sampling from the vector *π* of initialization probabilities for states {0, 1, 2}. Specifically, *π* is a vector of length 3 and which we initialize either to uniform or based on prior knowledge, and update at each step. Between each adjacent pair of states, we set the probability of transitioning between state *t*_1_ and state *t*_2_ with 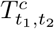. We parametrize *T*^*c*^, with a species-specific estimate of transition probability *p*_*c*_ such that *E*(transitions) = *M*_*c*_*p*_*c*_. Using our estimate for *p*_*c*_, we can define an idealized 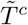:

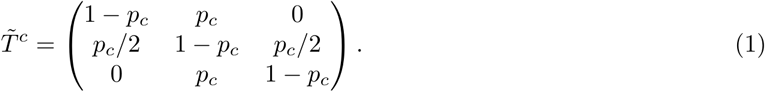

In practice, to account for rare genotype transitions and measurement errors, we add an entropy term *ϵ* and use the row-normalized matrix 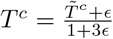. In cases where we have simulated data such as that described in the next section, we can initialize *T* by counting transitions for representative datasets (see Section 4.1.2). Finally, in the simpler case where there is no breeding structure, we can simply set each row of *T*^*c*^ to uniform. This is equivalent to assuming no relationship between SNPs, like in other models.

We now tie the chains back to the collected data. Using the collected matrices *A*^*c*^ and *D*^*c*^, we take the probability of seeing 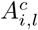 variant counts with depth 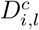 as Binomial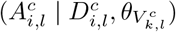, where we assume that the inferred cluster of cell *i* is *k* and the inferred genotype is 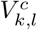. *θ* is a vector of variant allele frequencies for each state. We initialize *θ* to [0, 0.5, 1], but infer it with a Beta(*θ*_*t*_ | *α*_*t*_, *β*_*t*_) prior to allow for real-life behavior differences. The clustering *Z* is a vector, where *Z*_*i*_ is an integer representing the maximum likelihood clustering for cell *i*. We initialize it with a Categorical(*Z*_*i*_|*κ*) prior. Table 1 summarizes these governing distributions.

**Table 1.**
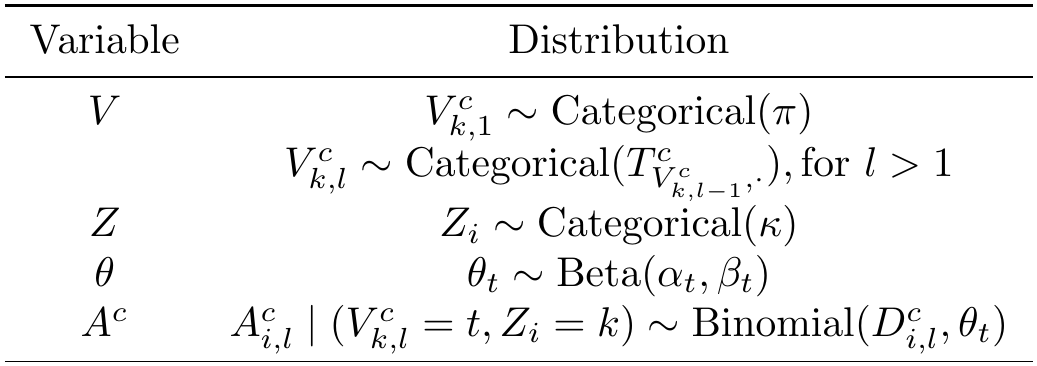
A summary of governing probability distributions for each model variable.

| Variable | Distribution |
|---|---|
| $V$ | $V_{k,1}^{c} \sim \mathrm{Categorical}(\pi)$ |
|  | $V_{k,l}^{c} \sim \mathrm{Categorical}(T^{c}_{V^{c}_{k,l-1},\cdot}), \mathrm{for}\ l > 1$ |
| $Z$ | $Z_{i} \sim \mathrm{Categorical}(\kappa)$ |
| $\theta$ | $\theta_{t} \sim \mathrm{Beta}(\alpha_{t}, \beta_{t})$ |
| $A^{c}$ | $A_{i,l}^{c} \mid (V_{k,l}^{c} = t, Z_{i} = k) \sim \mathrm{Binomial}(D_{i,l}^{c}, \theta_{t})$ |

We now summarize the model we have presented in two ways. First, we represent it graphically with Fig. 2B. There, the graph structure shows the previously defined relationships between variables, where the arrows indicate the dependencies. We pair this representation with the log joint probability of the model, which we present below. Note that we work in log-space for improved numerical stability and ease of computation.

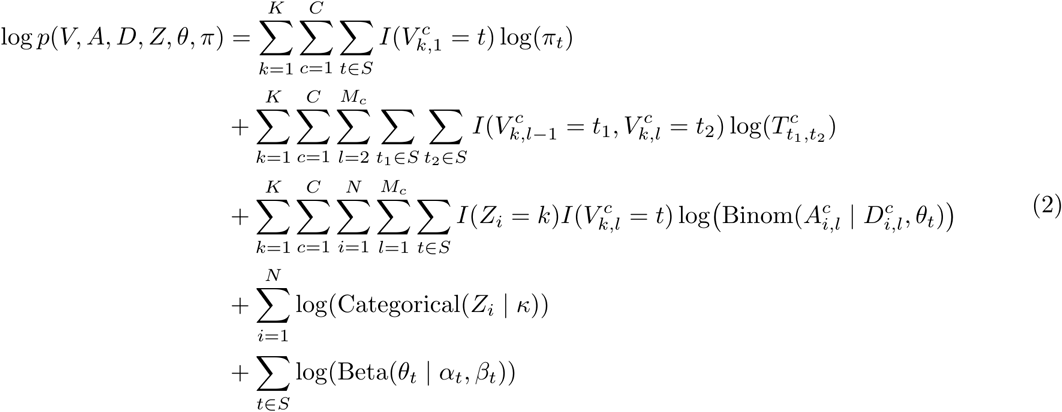

To optimize the model, we perform EM iterations until convergence, the update steps of which derive from the log-joint probability. We present details on these iterations, as well as the initializations of the variables, in Section 4.1.

In the applications we discuss below, we take a number of further practical steps to run the model optimally. Notably, for datasets with doublets, we created a doublet detection strategy which we detail in Sec. 4.4. Doublet detection allows us to filter out doublets which would otherwise hurt clustering and downstream applications. Additionally, we re-initialize and re-run the model multiple times to avoid local minima. For the final result, we choose to create a consensus of all the clusterings across restarts. This allows us to create a stronger final clustering instead of just discarding unused EM restarts (Sec. 4.3).

In the following subsections, we give practical demonstrations of DemuxHMM on both real and simulated data. We first demonstrate that our method is practical with the Diversity Outbreeding (DO) framework for mice using an existing real dataset (Sec. 2.2). Then, we move to a harder case of Drosophila embryos in simulated datasets (Sec. 2.3). The latter allows us to benchmark our performance in clustering and doublet detection against existing methods. Additionally, we can test a range of experimental conditions, to see where the method is most useful.

### 2.2 DemuxHMM Achieves Strong Performance on Pooled Diversity Outbred Mouse Data

We next sought to test DemuxHMM on the Diversity Outbred (DO) mouse line [27]. The DO approach differs somewhat from our simple example in Fig. 1 in that it uses 8 genetically diverse founder strains to create a large breeding pool based on Collaborative Cross descendants of the founders. However, that choice creates greater diversity per generation and is fully compatible with our HMM design. The specific DO mouse dataset we used came from Dillard et al. [22]. The dataset featured bone-marrow-derived stromal cells from 77 mice. After our variant calling and filtering (Sec. 4.5), there were 86,971 cells and 23,252 variants. Notably, the experiment pooled mice into batches of 4 or 5, giving us a partial ground truth. In our tests, we held out the pool labels and treated the whole dataset as one large pool, as would be the case in a massive-scale dataset.

Since the pool labels are only a partial ground truth, we evaluated demultiplexing using clustering homogeneity [28], a metric that can accommodate partial labels. Roughly speaking, homogeneity is an information-theoretic measure of how pure our clustering is in terms of pool labels. One should expect a good cluster to only have cells from one pool, while a bad cluster may have cells from multiple pools. Suppose *L* = {*q* | *q* = 1…*Q*}, are some class labels, in this case our ground-truth pools. Then, for our clusters *k* = 1…*K*, let *C* = [*c*_*q,k*_] be a contingency table. Here *c*_*q,k*_ represents the number of cells with pool label *q* that we assign to cluster *k*, where there are *N* total cells. As given by Rosenberg and Hirschberg [28] in the non-degenerate case, we define homogeneity, *h*, as:

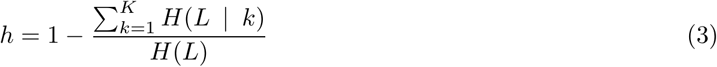

where *H*(*L*) and *H*(*L*|*k*) are entropy and relative entropy:

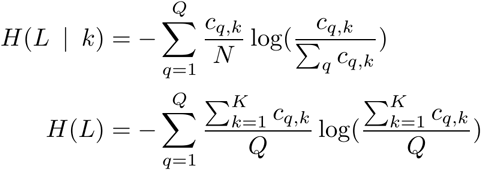

The notion of homogeneity allows us some measure of demultiplexing evaluation despite only having a partial ground truth.

With regard to DemuxHMM parameter selection, the main challenge stemmed from estimating transition matrices *T*^*c*^. In Sec. 4.6 we detail how estimates of per-generation recombination rates in DO mice [27] allowed us to compute transition probabilities, *p*_*c*_, and hence matrices *T*^*c*^, as well as other parameter choices. Another parameter question that may occur to the reader is how to choose the number of clusters, *K*. Here, we did have a ground truth and so used *K* = 77, but this is not a given across all datasets. To answer the question, we detail a heuristic method based on the model objective converging to a maximum as we reached the true number of clusters (Sec. 4.7). We note that this approach is not computationally feasible with all competing methods, and as such we did not use it for the tests that follow.

For the actual test of demultiplexing, we compared DemuxHMM to Vireo and Souporcell3 on the full dataset. Since the dataset contained doublets, we opted for a two-pass approach. That is, the first pass fitted each model and detected doublets, while the second pass refit each model after filtering out detected doublet cells. In the two-pass test, DemuxHMM scored *h* = 0.974 *±*0.003, Vireo scored *h* = 0.928 *±*0.007 and Souporcell3 scored *h* = 0.946*±* 0.006. For DemuxHMM and Vireo we reported these results as a mean of 5 repeats alongside their standard deviation. For Souporcell3, we only used 3 repeats due to the lengthy run time. DemuxHMM detected 16,805 *±*306 doublets, Vireo detected 15,528*±* 522 doublets, and Souporcell3 detected 13,126*±* 1,267. From these results, we saw that DemuxHMM performed noticeably better than the other methods, while marking slightly more cells as doublets. All three methods had very low variance in performance for this test. In Fig. 3B, a plot of per-cluster error makes the performance difference even clearer. We define per-cluster error as 1 *− h*_*k*_, where 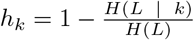. In this view, DemuxHMM had fewer error-prone clusters, with only 9% of clusters having over 1% error. For Souporcell3, this rose to 12% of clusters, and for Vireo, 19%. On the higher end of error, only 6% of DemuxHMM clusters exceeded 20% error. This was in contrast to 9% for Souporcell3, and 13% for Vireo.

**Fig 3.**
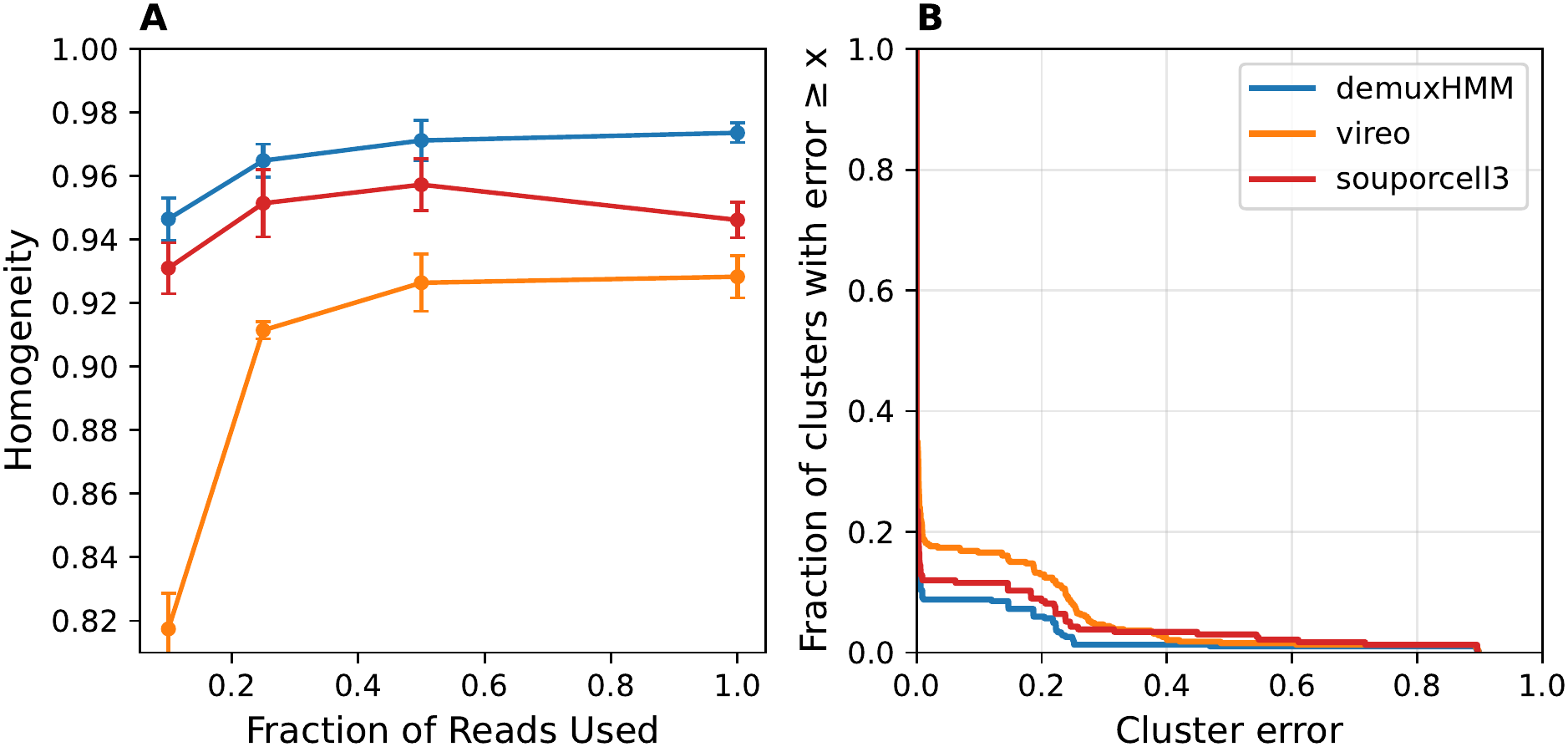
A comparison of demultiplexing performance on the DO mouse dataset. Panel **A)** shows two-pass homogeneity scores for the demultiplexing methods on the DO mouse dataset for different levels of reads downsampling. Points are a mean of 5 repeats for DemuxHMM, and Vireo. For Souporcell3, the points use 3 repeats due to run time. Error bars represent standard deviation over resampling repeats for fractions less than 1 and over full methods reruns for the non-downsampled data. **B)** A cumulative error plot of homogeneity for each method. The lines indicate the percentage of clusters with an error above the amount given by the x-axis. The plot includes all clusters across all repeats.

In terms of run time, we noted that while DemuxHMM finished in 5.25 *±*0.43 hours and Vireo finished in 11.08*±*2.60 hours, Souporcell3 took 149.16*±*13.50 hours despite our giving it 60 CPU threads. We ran these tests on a system with a Nvidia RTX 4500 Ada Generation GPU, and an AMD EPYC 9654 CPU. Practically, this means Souporcell3 may be challenging to run on such large datasets for groups without access to high-performance computing. In contrast, one can run DemuxHMM on a consumer-grade GPU with *≈*24 GB of VRAM in just a fraction of the time. In our experience, there is also room to downsample the SNPs with minimal impact on performance, letting the model fit on GPUs with less than 16 GB of VRAM for this dataset.

Our next objective was to determine the effect of downsampling variant counts on demultiplexing performance. To do this, we downsampled the counts to 10%, 25% and 50% of the original and re-ran demultiplexing (see Sec. 4.5 for the procedure). We ran DemuxHMM and Vireo with 5 resampling repeats for bootstrapping, and Souporcell3 with only 3 repeats due to computational cost. Fig. 3A shows the results of the test. There, we saw that DemuxHMM maintained a clear performance advantage across all downsampling levels. Surprisingly, we also noted that Souporcell3 actually performed better than with the full dataset, particularly when we used 50% or 25% of the reads. We hypothesize that Souporcell3 is sensitive to low-information, high-depth SNPs. However, neither DemuxHMM nor Vireo displayed the same level of sensitivity. On the low-depth end of the spectrum, at the 10% retention level, DemuxHMM’s advantage widened significantly, especially compared to Vireo. This gap suggests that the HMM structure becomes more critical at shallower sequencing depths. Therefore, using our HMM-based approach would provide a strong advantage in shallow-depth studies.

We next sought to examine clustering error more finely, beyond global clustering agreement metrics. In Fig. 4, we viewed each cluster on the full dataset as a pi chart colored by the ground truth pools of its cells. We assigned each cluster to a pool based on the majority of cells. We compared this against Souporcell3 (we included the same view for Vireo as a supplement). Note that we used the best repeat run for the respective methods. On inspection, we noticed that DemuxHMM assigned the correct number of clusters to a pool more often. We consider a cluster as part of a pool if the largest share of cells in that cluster belongs to the pool. DemuxHMM assigned the correct number of clusters for 12 of 16 pools, while Souporcell3 only got the number correct for 8 of 16 pools. For both methods, we noticed that some clusters split into multiple pool segments. DemuxHMM showed 5 clusters with a non-negligible amount of cells from the wrong pool, while Souporcell3 showed 11. We hypothesize that these splits occurred due to some difficult-to-assign cells that failed to map to their own cluster. Depending on the goals of the group working with the data, it would be reasonable to filter out these clusters. Finally, we noticed that for the fifth pool from the top, Souporcell3 had a cluster with 3,315 cells which cannot all be accounted for in the DemuxHMM clusters for that pool. Upon inspection of the cluster, we found that DemuxHMM had marked the majority of the cells as doublets. We suspect that these are indeed doublets because this cluster was an outlier in terms of the number of cells.

**Fig 4.**
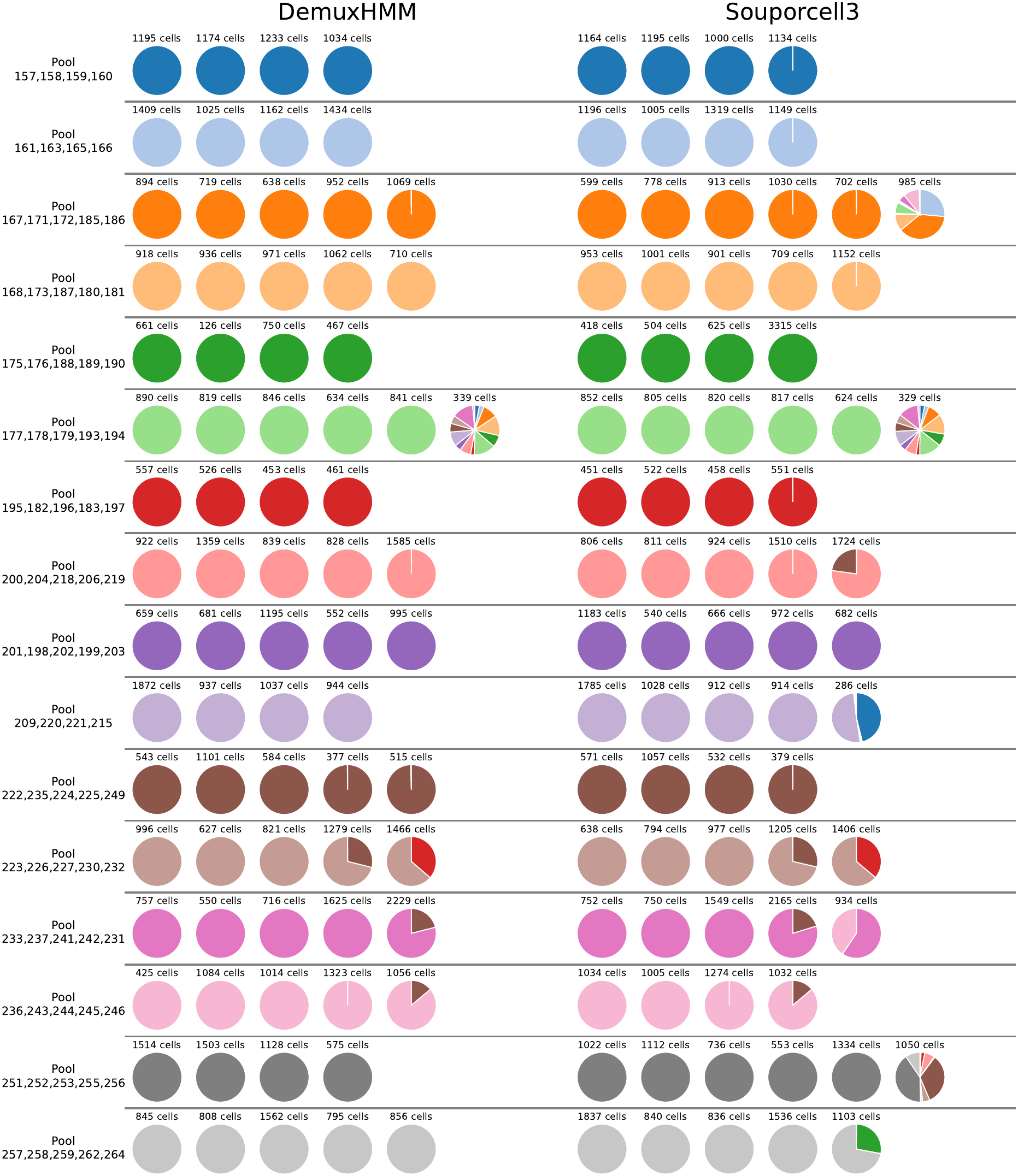
A view of clustering on the DO mouse dataset comparing DemuxHMM and Souporcell3. Each cluster is represented as a pi chart and annotated above by the number of cells contained within. The colors of the pi chart represent the cluster’s composition with regard to ground truth pool identity. Clusters are assigned a row based on majority pool fraction. Pool names represent the lab-assigned numbers for mice sequenced in that pool. Clusters for both DemuxHMM and Souporcell3 shown are the result of two-pass demultiplexing and are filtered for doublets.

Overall, we saw that DemuxHMM achieved strong performance in terms of homogeneity score on the Dillard et al. dataset. This dataset demonstrated a clear advantage of the HMM approach for a real-world breeding scenario. Additionally, the dataset is large in terms of both cell counts and individual counts, showing that we can handle both in quantity while maintaining good performance and run time. In Sec. 2.3 these performance trends continue on simulated data, including rigorous benchmarks of potential real-world failure modes (low levels of breeding, low depth, doublets, etc…).

### 2.3 DemuxHMM Leverages Recombination for Better Demultiplexing on Simulated Fly Data

In this section, we tested DemuxHMM against existing self-genotyping methods on synthetic datasets with our proposed recombination barcoding method. Through a series of tests, our goal was to establish that DemuxHMM performed better than existing methods on datasets with a breeding structure and that the HMM was the driver of this performance. We then established that the performance levels we observed are usable for downstream analysis. Finally, we showed that we could achieve good performance over a range of experimental conditions (breeding generations, depth, available SNPs, and the presence of doublets) rather than a few highly tuned settings.

To perform the aforementioned tests, we created simulated datasets based on *Drosophila melanogaster*. The datasets simulated sibling mating of *G* generations of two genetically diverse founder strains from the Drosophila Synthetic Population Resource (DSPR) [29]. This simulation provided a flexible test environment, allowing us to test DemuxHMM under a variety of realistic conditions, including the presence of doublets. Compared to the DO mouse dataset, we had a full ground truth, giving a more informative performance comparison, and allowing us to evaluate doublet detection accuracy. Overall, we presented results showing that our method achieved high performance, with low runtime, even for large numbers of individuals, across a range of experimental settings.

We chose *Drosophila melanogaster* as our model organism because it provided a realistic and conservative benchmark for our joint recombination barcoding and demultiplexing approach. *Drosophila* has four chromosomes. Out of these, chromosome 4 is small and non-recombining, meaning it contributes too little to diversity to be useful for our demultiplexing approach. As a result, our model needed to achieve sufficient barcode complexity with only three usable chromosomes. This is in contrast to other species, such as mouse, where the presence of many recombining chromosomes may make it easier to achieve high barcode complexity. At the same time, *Drosophila* offered practical advantages for evaluating the DemuxHMM framework. In the preferred scenario for breeding, experimentalists would use two highly divergent parental strains. *Drosophila* can have high levels of variation between strains [30], with the DSPR A6 and the DSPR A4 strains having a SNP approximately every 204 base pairs [31]. Accordingly, we chose these as the parental strains for our simulation. Furthermore, Fiston-Lavier et al. modeled recombination rates by chromosome location for *Drosophila* [32], allowing us to realistically simulate recombination, including rare double crossover events that can occur during meiosis. Taken together, these factors suggest that the results with simulated data should closely reflect those we see in a real experiment.

We briefly outline our simulation procedure for all synthetic datasets. We provide an in-depth description in Section 4.8. The procedure begins with a SNP profile for a DSPR A4 father relative to a DSPR A6 mother. We generated offspring by modeling meiosis and choosing recombination points according to modeled recombination rates. We note that in *Drosophila*, males do not generally experience recombination, and overall there are only three recombining chromosomes. Thus, all else equal, other species may require fewer breeding generations to achieve similar performance. The simulation created a pool from which we selected breeding pairs to simulate the next generation. After *G* generations, we sampled cell profiles from a real head and body scRNA-seq dataset from the Fly Cell Atlas [33]. Using these cell profiles, combined with our simulated chromosomes, we sampled our variant and depth matrices (*A*^*c*^ and *D*^*c*^, respectively). To improve demultiplexing performance, we filtered to cells with variant counts on each of the recombining chromosomes (X, 2, 3). By following this general procedure, we generated datasets for a variety of possible experimental conditions.

Our primary metric for measuring clustering agreement was the Adjusted Rand Index (ARI) [34]. ARI is a measure of clustering agreement adjusted for chance. It measures how close a clustering is to the true partitioning and is a common metric for evaluating clustering performance. More precisely, in [35], William Rand defined the Rand Index as:

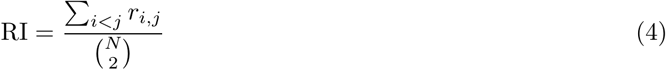

where *r*_*i,j*_ = 1 if either cell *i* and cell *j* are in the same cluster in both the reference partition and evaluation partition, or they are in different clusters to each other in both the reference and evaluation partition (sometimes referred to as a true positive and true negative). Otherwise, *r*_*i,j*_ = 0 (false positive or false negative). The Adjusted Rand Index then applies chance-correction to the score:

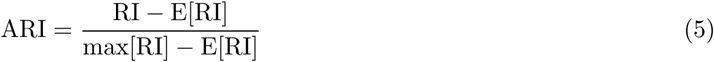

where max[RI] = 1 for our use case. While ARI is not the only way to measure demultiplexing performance, it is the most straightforward metric to compare across methods.

We first simulated a scenario where the total number of cells was fixed, but the number of individuals varied, modeling a typical sequencing situation. In this scenario, a researcher expects to capture a certain number of cells at an acceptable depth, constrained by their budget and the current single-cell technology. The problem is then to determine how many individuals to divide the cell budget between. With a limited cell budget, allocating too few cells per individual can result in noise dominating the signal, affecting demultiplexing performance and downstream analysis. Thus, we wished to determine the trade-off that occurred as we increased the number of individuals. Using 20,000 total cells at 10,000 UMI per cell, and 22 generations of breeding, we varied the number of individuals from 10 to 500 to create simulated datasets. Approximately 75% of embryos and 50% of cells survived the filtering process (Fig. 5C). We chose 22 generations of breeding based on performance tests, but noted that performance remained robust to lower numbers of breeding generations (see later tests in this section). DemuxHMM outperformed the competing methods across all levels of individuals in terms of ARI (Fig. 5A). We then repeated the same experiment for DemuxHMM and scSplit at a lower depth of 2,500 UMI. At this depth, the performance gap between the two widened significantly, with DemuxHMM retaining strong performance.

**Fig 5.**
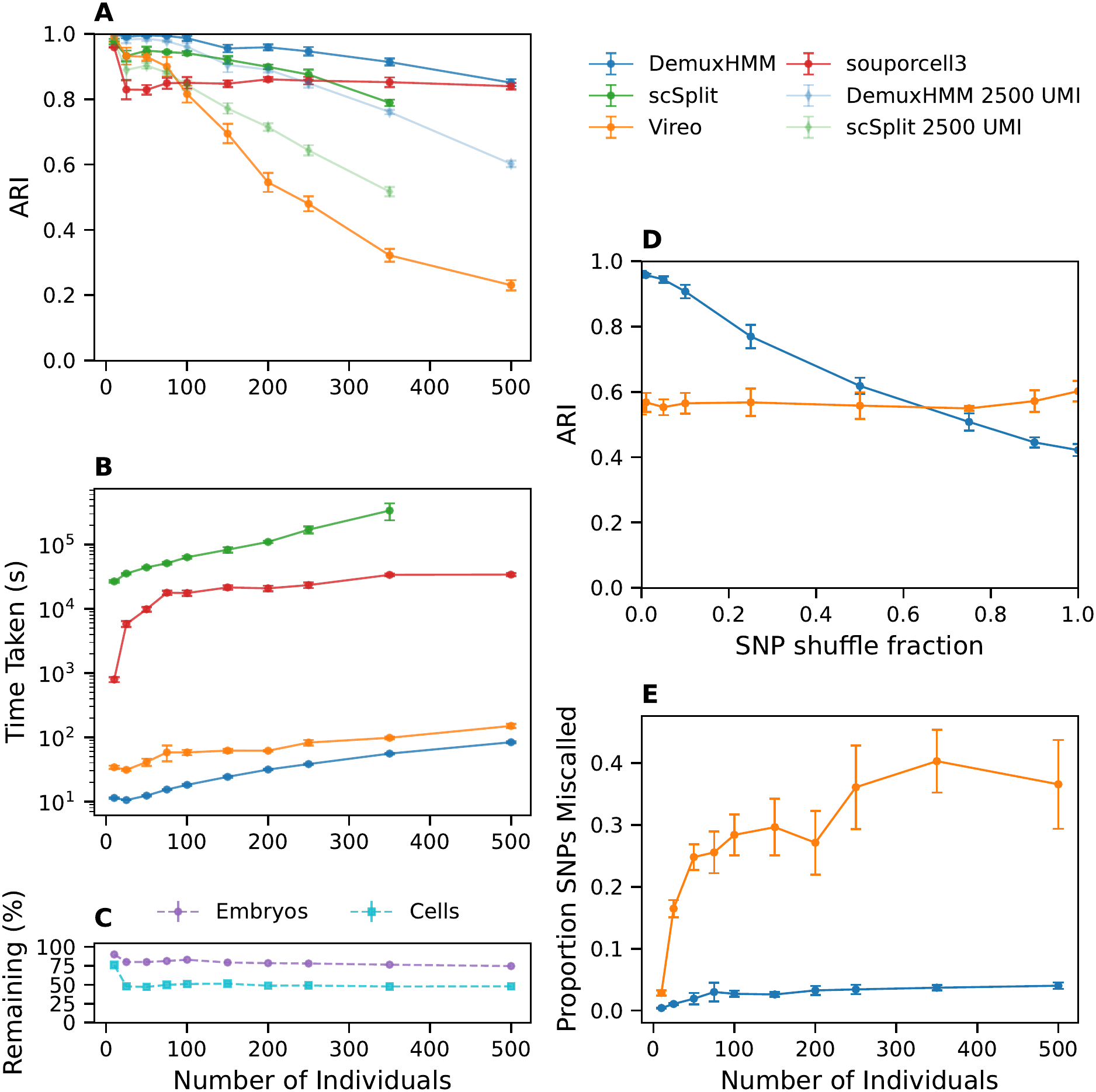
A comparison of fixed-budget demultiplexing performance as the number of individuals is varied. Each dataset has a fixed total cell budget of 20,000 cells pre-filtering. The datasets have 22 generations of breeding, and either 10,000 UMI (opaque points) or 2,500 UMI (transparent points). All results are taken as a mean of 6 trials across dataset resamplings. Error bars indicate standard deviation. Runs with 500 individuals for scSplit did not finish within 7 days of runtime and were not included. **(A)** The demultiplexing performance as measured by the Adjusted Rand Index (ARI). **(B)** A comparison of method run times over the 10,000 UMI datasets. The y-axis is time in seconds, presented on a log scale. **(C)** The number of cells and embryos remaining after filtering. **(D)** A performance comparison between DemuxHMM and Vireo with 200 individuals as an increasing fraction of SNP positions is shuffled. **(E)** Shows genotyping error as the proportion of incorrectly called SNPs taken as a mean across individuals. The sweep uses the same datasets seen in panel A. As before, the points themselves are a mean across data resamplings and error bars are standard deviation.

A natural follow-up question was whether the addition of the HMM truly drove the performance difference. We addressed this question through the following empirical test: suppose we were to incrementally break the advantage of the HMM by removing the ordering and chromosome information associated with the SNPs. We simulated this test by moving some fraction of the SNPs onto a fake chromosome ‘Z’ which did not preserve any genomic ordering. For the shuffled chromosome Z, DemuxHMM used a uniform transition matrix. If the HMM aspect is driving performance, the greater the fraction of SNPs shuffled, the worse the performance should be for DemuxHMM. On the other hand, non-HMM methods such as Vireo should remain relatively unaffected. Indeed, running this test on the 200-individual datasets yielded decreasing performance as we shuffled more SNPs, while Vireo (the method most similar to ours) kept a relatively flat performance curve (Fig. 5D). One might note that our method did converge to a lower performance than Vireo in this test. The gap may stem from implementation differences such as Vireo using variational inference rather than EM. Regardless, we saw the expected behavior as we shuffled SNPs, giving us some assurance that the HMM is driving the stronger performance.

In addition to testing performance, we compared run times between methods. We found that on a system with an Nvidia RTX 4500 Ada Generation, and an AMD EPYC 9654 CPU, DemuxHMM achieved run times orders of magnitude faster than scSplit and Souporcell3, and faster than Vireo (Fig. 5B). In particular, we saw that scSplit was unlikely to be scalable to larger real-world datasets despite having the second best performance after DemuxHMM. To further demonstrate the scalability of DemuxHMM, we generated three very large datasets with 1,000 individuals each (mean 757 retained), 250 cells per individual pre-filtering, and 22 generations of breeding. DemuxHMM demultiplexed these datasets with a mean ARI of 0.778*±* 0.013 and a mean run time of 2.23*±* 0.05 hours, demonstrating that the method scales to thousands of individuals while maintaining strong performance.

In the tests so far, we had focused on clustering quality metrics. However, applications such as crossover point mapping may be more dependent on the quality of inferred genotypes for each individual. Since we had ground truth genotypes for the simulated flies, we evaluated genotyping performance on the same fixed-cell dataset. We measured genotyping error as the proportion of SNPs we called incorrectly in an individual, and reported the mean across all individuals. In Fig. 5E, we saw that genotyping error was less than *≈*4% ( *≈*49 of 1,233 SNPs mis-called) for all levels of individuals. In contrast, Vireo saw rates above 25% for the majority of individual counts. With this result, we could reasonably conclude that DemuxHMM maintained acceptably low genotyping error across the full spectrum of conditions we tested.

Considering the performance levels we observed, our next question was whether that accuracy was enough for reasonable downstream analysis. To examine this question, we aimed to gain an intuitive understanding of the impact of worsening ARI levels on downstream analysis such as trajectory inference. We performed a simulated experiment with a time-course dataset of sea urchin embryonic development [36]. To simulate demultiplexing error, we randomly shuffled a percentage of cells between timepoints (Section 4.10). We did this several times for different percentages of cells to simulate different demultiplexing ARI levels. We performed trajectory inference on each dataset, quantifying the difference relative to the original dataset through an Earth Mover’s distance. Cell fate inference error increased roughly linearly with worsening ARI (Fig. 6). For selected cell types, fate plots remained qualitatively similar to baseline for an ARI of *≈*0.8. This suggested that the ARI levels DemuxHMM achieved in challenging regimes with high numbers of individuals were likely to be sufficient to preserve biologically meaningful trajectories.

**Fig 6.**
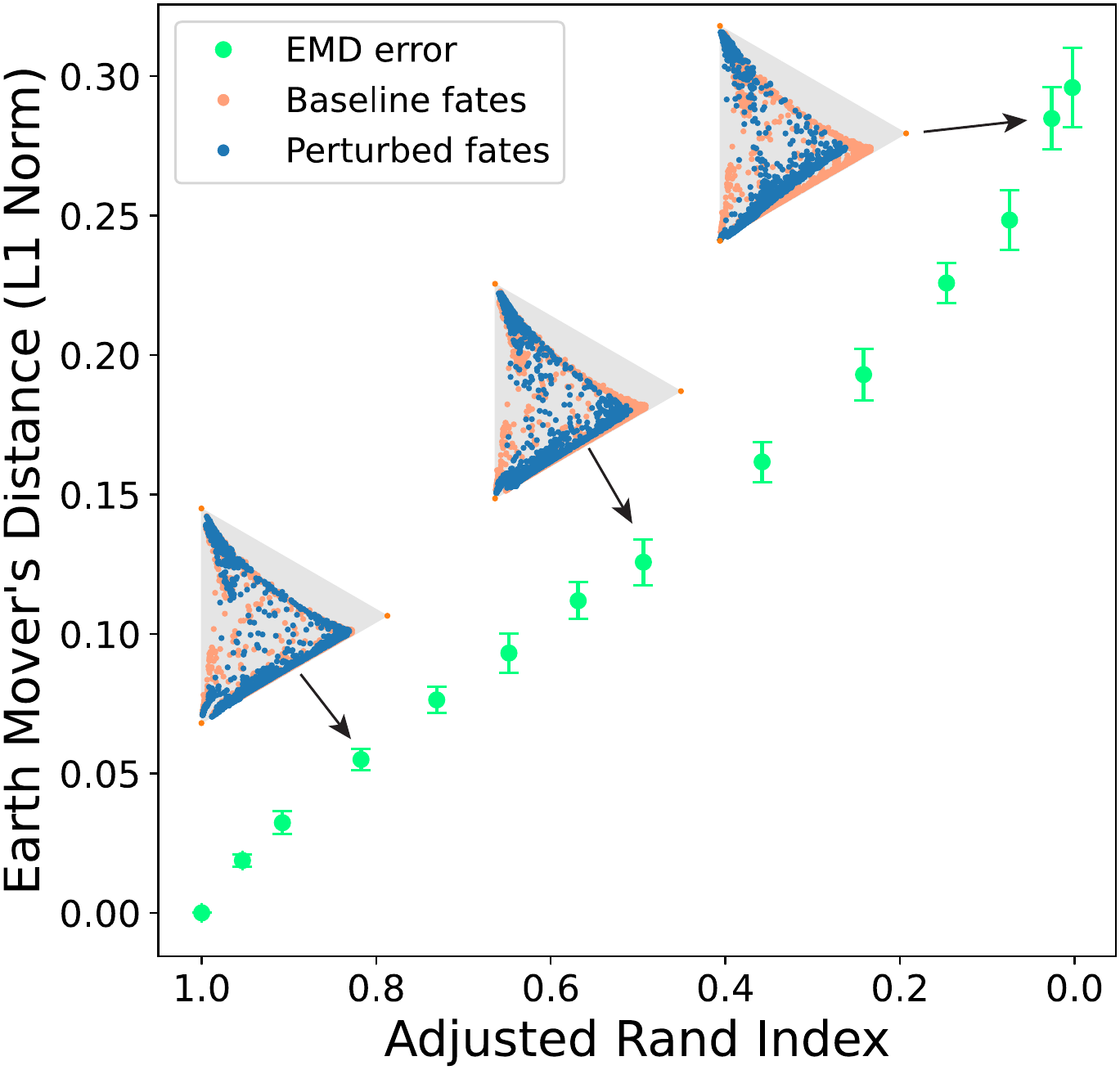
The error of inference on a *Lytechinus variegatus* (sea urchin) time series [36] as ARI changes. Green dots represent mean errors at a given ARI over 10 re-assignments. Error bars represent the standard deviation across these repeats. For selected dots, a 2D representation of probabilities for cells at 10 hours post-fertilization (hpf) to become Smooth Muscle Cells (SMC) versus endoderm cells versus other cell types are shown on a triangle plot. In those plots, orange dots represent baseline fates, whereas blue dots represent fates in the perturbed datasets. Datasets were perturbed by randomly shuffling time labels of cells in the dataset, with a changing percentage of cells being shuffled. ARI was then measured against starting labels. Fate matrices were then computed for each level, including baseline, as described in the original study. Error was measured using Earth Mover’s Distance (EMD) on the matrices for 10 hpf. The cost matrix used for EMD was made by taking the *L*^1^ distance between matrix rows (cell fate vectors).

Having seen the performance results, we needed to know whether we could reliably produce them over a wide range of experimental conditions. If it is too difficult to produce viable datasets, research groups may hesitate to create them. Accordingly, our goal was to give some guidance about suitable ranges of conditions (here, applied to Drosophila).

First, we evaluated the robustness of DemuxHMM over a range of breeding experiment conditions. Specifically, we performed a grid sweep over the number of breeding generations *G* and mean UMI counts per cell. This sweep simulated scenarios where we performed less than optimal levels of breeding, or sequencing depth was lower. In Fig. 7A, we saw that DemuxHMM maintained strong demultiplexing performance across a large band in this range, including at a very reasonable 2,500 UMI per cell for many levels of breeding generations. Thus, we can have some assurance that performance is possible for many reasonable depths and breeding generations.

**Fig 7.**
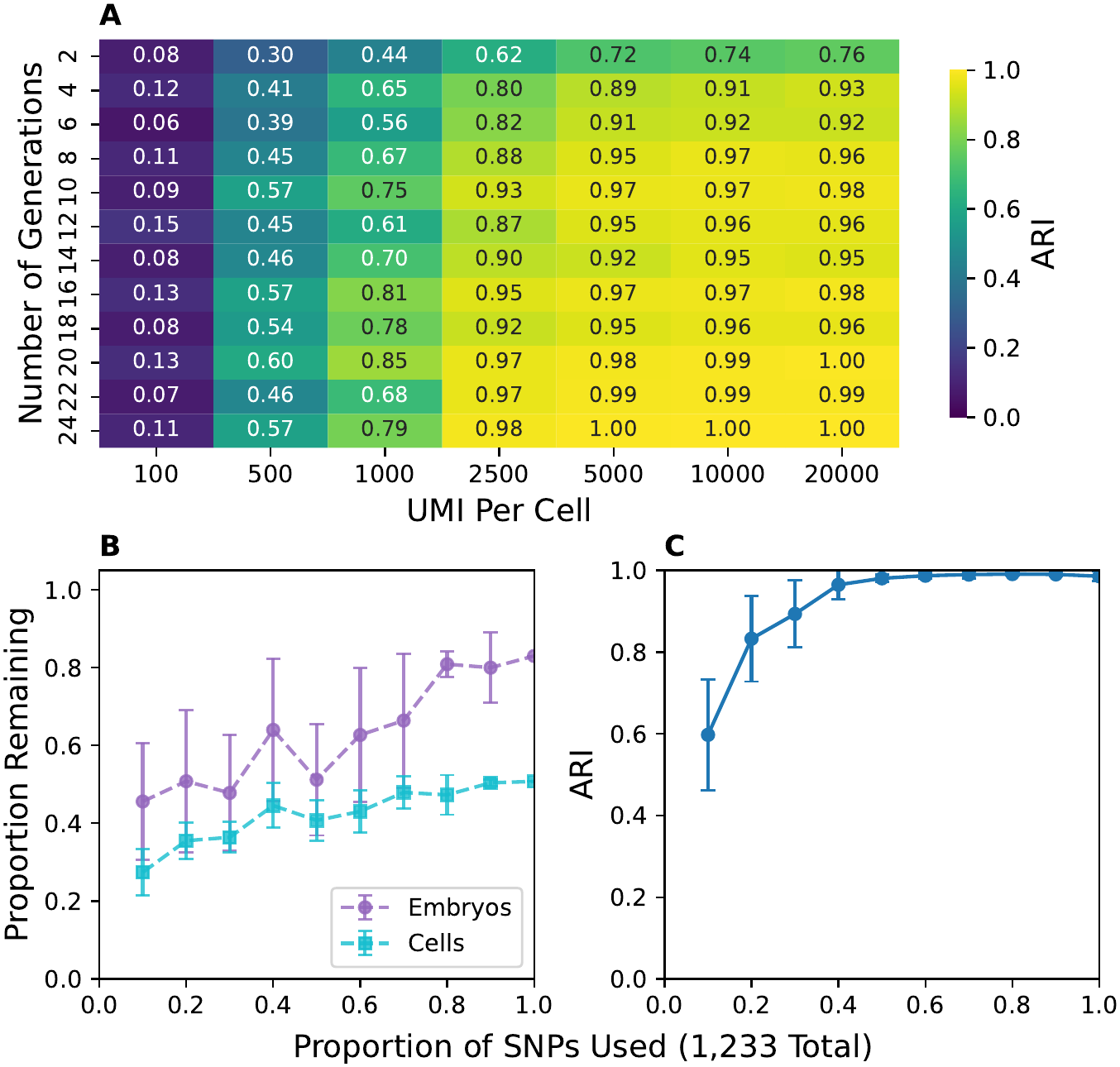
DemuxHMM performance (ARI) on simulated *Drosophila* datasets under changing experimental conditions. The datasets used have 100 individuals, and 250 cells (pre-filter) per individual. **(A)** A grid of DemuxHMM performance on datasets generated with changing numbers of breeding generations and mean UMI counts per cell. Values are a mean of 6 resampling repeats. **(B)** shows the proportion of embryos and cells that pass filtering as we downsample the number of usable SNPs. **(C)** shows performance as this occurs. For these panels, values are a mean over 10 data resamplings. Error bars indicate standard deviation.

Next, we sought to test how the genetic diversity of the parents (the total number of SNPs we saw) affected the accuracy of DemuxHMM. Specifically, we tested the sensitivity of DemuxHMM to the number of SNPs we observed. The ability to operate with fewer informative variants provides greater flexibility in the choice of parental strains. In the strains we analyzed, there were originally 1,233 SNPs retained after our initial filtering step. Starting with a dataset of 100 embryos with strong baseline performance (250 cells per individual and 10,000 UMI per cell), we downsampled SNPs at various levels. Performance remained largely unchanged until we retained approximately 40% of the original 1,233 SNPs (Fig. 7C). We noted that this is already a low number of SNPs compared to our DO mouse dataset, and as such we considered this to be a difficult test to start with. Aside from performance, the fraction of embryos passing quality filtering decreased approximately linearly with SNP downsampling, with 64% of embryos retained at 40% SNPs (Fig. 7B). This meant that lower levels of SNPs also left somewhat fewer usable embryos. However, on the whole, the results indicated that DemuxHMM tolerated substantial reductions in SNP density, providing flexibility when variation is a challenge.

Finally, we considered the presence of doublets. Doublets occur when two droplets merge during sequencing, creating a fused cell profile [37]. We wanted to be able to detect and filter a range of doublet rates reliably to remain useful even on doublet-heavy datasets. While we were able to detect doublets in the DO mouse dataset, we lacked a doublet ground truth to evaluate detection performance. With the simulated dataset, we could thoroughly benchmark doublet detection. We generated datasets with 20,000 cells pre-quality-control, 50 individuals and 10,000 UMI per cell with doublet rates of 2.5%, 5%, 10%, and 20% (see Sec. 4.8.1 for details). We measured doublet detection quality through a number of metrics including precision, recall, area under the precision-recall curve, as well as ARI when we filtered detected doublets out (Fig. 8). At our selected doublet threshold (a probability of ≥ 90%), DemuxHMM had somewhat lower precision than scSplit and Souporcell3; however, it had the highest recall of all methods. Additionally, we had the highest area under the precision curve. This meant that our method had the most room to trade off between precision and recall depending on the doublet threshold we set. We also showed clustering ARI after doublet filtering. While pre-doublet-filtering ARI was moderate (see Supp. Fig. 10), our two-pass ARI was the best of all methods, giving a near-perfect clustering for all levels of doublets. Consistent with the DO mouse results, the higher recall and slightly reduced precision explained why we marked somewhat more cells as doublets. We likewise observed the two-pass ARI results in the DO-mouse data, as we saw that DemuxHMM had the highest two-pass homogeneity there. Overall, the doublet detection results showed that the method was effective in detecting and filtering doublets up to a very high rate of 20%.

**Fig 8.**
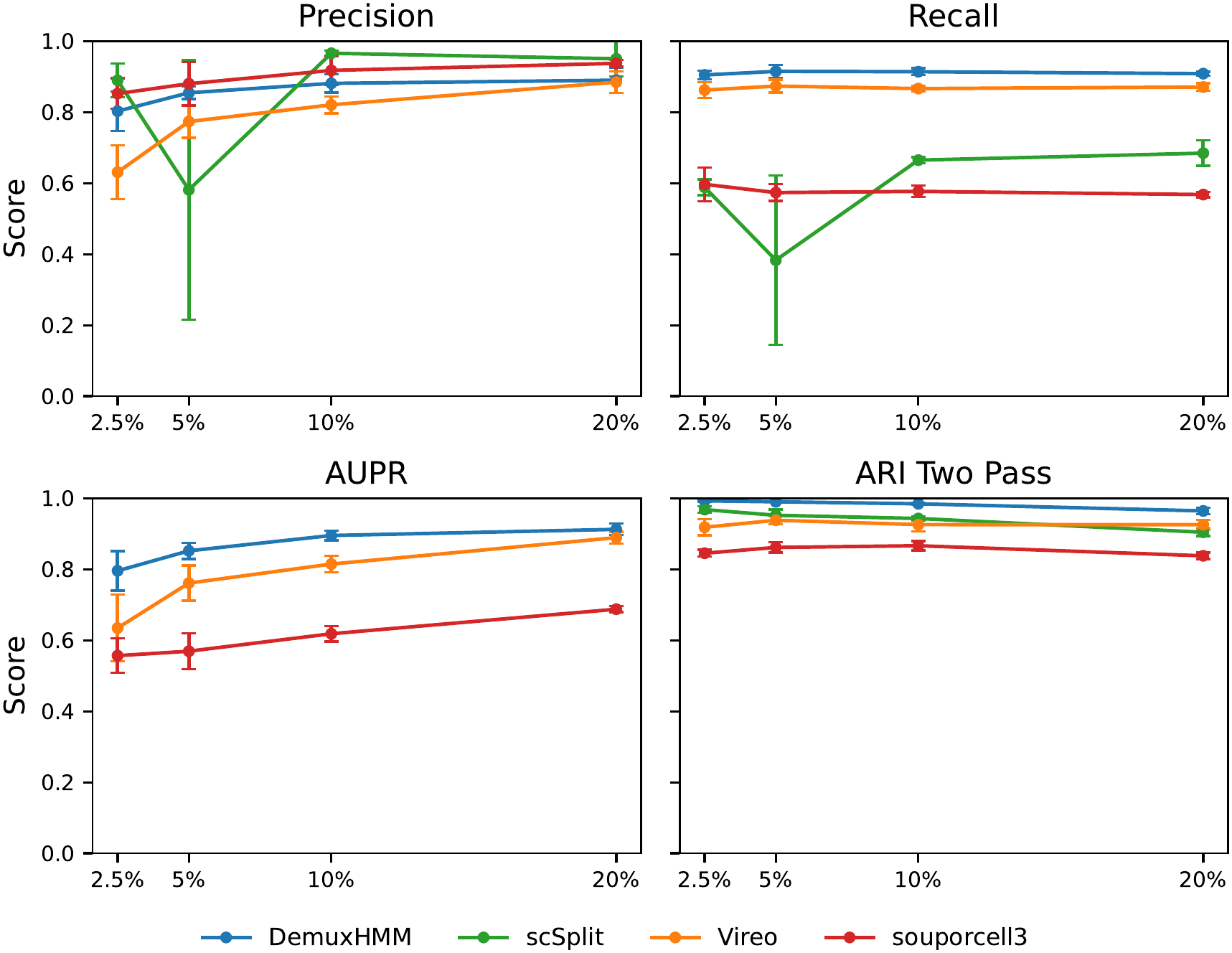
A benchmark of doublet detection performance across demultiplexing methods. Performance is evaluated on simulated datasets with varying doublet rates containing 20,000 cells and 50 individuals (pre-quality-control). Evaluations from left to right, top to bottom are precision, recall, area under the precision curve (AUPR), and two-pass ARI (where we filter doublet cells before demultiplexing). Values given are the mean of 6 data resamplings. Error bars indicate standard deviation. For each plot, the x-axis is the doublet rate, and the y-axis is the relevant metric score. For each metric, 0 is worst and 1 is best.

Overall, our simulated results provided practical guidance on the setup of demultiplexing experiments. In the results, we showed that DemuxHMM maintained strong performance up to a high number of individuals, even when we had few cells per individual. The performance levels we achieved were also at a useful level for downstream analysis through trajectory inference. We saw that we could achieve this level of performance throughout a range of experimental conditions. In the presence of doublets, DemuxHMM had the best recall at the expense of precision, with room to trade precision for recall.

## 3 Discussion

In this paper, we propose a highly scalable strategy for creating large individual-resolved datasets by demultiplexing: DemuxHMM. DemuxHMM uses a Hidden Markov Model (HMM) to leverage simple breeding schemes for recombination-based barcoding. The HMM approach allows the method to utilize an underlying structure for variant-based demultiplexing, giving significant performance improvements over existing methods. This improved performance allows us to demultiplex higher numbers of individuals effectively, to the scale required for high-density time courses. We showed that our method works with real mouse data using the Diversity Outbreeding (DO) scheme. Additionally, our results demonstrated that the method could demultiplex hundreds to thousands of individuals in a simulated Drosophila dataset. This performance held through a range of experimental conditions (breeding generations, UMI counts per cell and the presence of doublets). Looking forward, we expect advances in single-cell technologies to further enhance performance, allowing even greater scalability. These combined results place our method in a unique position to be effective in demultiplexing new, large, structured datasets.

With the release of our approach, there are now a number of exciting opportunities to create test datasets. First, we already showed that our approach works with the DO scheme. This opens the door to creating larger DO mouse datasets for a variety of applications. With regard to other species, our strong results with the simulated data suggest another obvious step would be to create a real *Drosophila melanogaster* dataset for demultiplexing. Finally, creating a large time course on the scale of hundreds of individuals appears possible. Such a time course would require modifications to the breeding approach to allow for fully asynchronous preparation of generations of individuals. However, this time course could provide significant gains to trajectory inference studies of development.

Aside from time series trajectory inference, the approach of demultiplexing by genotyping hundreds to thousands of pooled recombinants from single-cell datasets has broad applicability. Because this approach simultaneously provides genotype and gene expression information for large numbers of individuals, we anticipate several other avenues of application. Instead of embryos of different developmental stages, when one pools embryos of the same stage or even tissue from different individuals, one can estimate expression variance not only for each cell type - needed for transcriptional kinetic inference [38] - but also between individuals. As a quantitative trait, researchers have used gene expression and its variance between individuals and populations to infer natural selection on expression differences [5]. Further, the reconstruction of the recombinant genotypes from the breeding scheme inherently requires the inference of crossover (CO) positions; given CO placements for enough individuals in the pool, one can construct genome-wide recombination maps, which traditionally required individually sequencing hundreds to thousands of recombinant offspring [7]. This extends to QTL trait mapping when researchers pool individuals with phenotypes of interest. Lastly, the combination of structured genotype and gene expression variance enables the detection of associations between gene expression differences and possible genetic determinants, i.e. eQTLs [6]. This obviates the need for RNA-sequencing and genotyping large numbers of samples, as one can accomplish both within one sample pool. As single-cell sequencing methodologies advance, the number of barcodes will likely increase and the sparsity of the data will decrease. Thus, we can expect the accuracy of genotyping and capacity for resolving individuals to improve over time, further widening the applicability and efficacy of DemuxHMM in these additional applications.

Still, challenges remain in verifying DemuxHMM in more challenging experimental scenarios, such as those involving strains with lower genetic variation. In this setting, one potential solution to capture more variation is to also perform variant calling for multi-nucleotide variations and indels, which would require no modification to the DemuxHMM framework to use. Additionally, there is evidence that other sequencing modalities such as scATAC-seq perform well in demultiplexing applications [39]. DemuxHMM should be able to use variant calls from these modalities with little modification.

Overall, as inference methods develop, the need for high-density time courses with hundreds or thousands of samples will rapidly grow across many model species. As datasets surpass dozens of distinguishable individuals, manual individual-level processing by a technician will become infeasible. At the same time, improvements in single-cell library preparations will enable the capture of increasing numbers of cells, and hence individuals, per experiment. These trends will amplify the need for demultiplexing methods that can scale to large numbers of individuals. Our simulations showed that DemuxHMM will have strong performance on such datasets when used in combination with a breeding scheme. Together, our work positions the combined experimental and computational framework of DemuxHMM as a foundation for creating large-scale single-cell time series without the need for individual-by-individual processing.

## 4 Materials and methods

### 4.1 Model Initialization and Update Steps

The DemuxHMM model is based on predicting a chain of SNP states for each chromosome of each individual. Cells are then assigned a maximum likelihood cluster (individual of origin) based on their own states. The model is best summarized by the graphical diagram in Fig. 2B. In this section, we will introduce the variables from this model and then provide workings for their respective EM update steps.

#### 4.1.1 *π* and *V*

*V* is the primary driver behind our inference and is a set of matrices for each chromosome *c*. Each 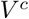 has a dimension of *K × M*_*c*_, where *K* is the total number of individuals and *M*_*c*_ is the number of variants on chromosome *c*. 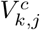 is a state in {0, 1, 2*}*, representing homozygous reference, heterozygous, or homozygous variant, respectively, for each variant position. 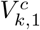 is initialized according to *π*, which is a vector of length 3 of initial probabilities for each state. *π* is dependent on species-specific recombination rates, site locations, and other factors. In practice, it has proven beneficial to use 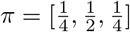 when no prior information is known, and then learn *π*. This initialization can be justified by the fact that two pairings of variants on the homologous chromosomes can lead to the heterozygous state, but only one pairing of variants can lead to each homozygous state. This initialization strategy was employed for our simulated dataset. After setting *π*, we sampled states using probabilities in *π* to initialize 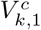. Further values 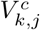 were initialized by sampling transitions from *T* (described below).

#### 4.1.2 T^c^

*T*^*c*^ represents the transition probabilities between variant states for chromosome *c*. We describe a theoretical approach in Section 2.1 that requires a guess on estimated transition counts. For the simulated data, we generated high-quality datasets with 1,000 individuals for a range of 1-25 breeding generations. We then counted the number of state transitions occurring and used those for initializations. Datasets for the described trials were generated independently and used this saved estimate to inform *T*.

#### 4.1.3 *Z*, and *κ*

*Z* represents the clustering of each cell and has dimension *N*, where *N* is the total number of cells. An upper bound on the number of clusters is assumed to be known as *K. Z*_*i*_ is a number in 1,…, *K*. We note that we observed that overestimating *K* is not problematic for accuracy, as extra clusters tend to be assigned no cells. However, there is a speed impact on the inference. *Z* is initialized according to its prior, *κ*, which is taken to be *Uniform*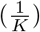.

#### 4.1.4 *θ, α*, and *β*

*θ* is a vector of dimension 3 specifying the probability of seeing a variant count for each genotype. In the ideal scenario, this is [0, 0.5, 1]. However, biological factors can affect alternate allele rate, so we can instead initialize *θ* using *Beta*(*α, β*), where *α* and *β* are priors. We used the values of *α* and *β* given in [18]. For the simulated dataset, we initialized *θ* with the learned values of [0.006, 0.469, 0.988].

#### 4.1.5 *A* and *D*

*A*^*c*^ is an *N × M*_*c*_ matrix counting the number of alternative allele counts seen at chromosome *c*, cell *i* and variant positions 1,…, *M*_*c*_. *D* has the same dimension and represents the total number of UMI counts seen at that position (depth). For real data, both are initialized from variant calling from cellsnp-lite [40]. However, other variant callers, such as a combination of FreeBayes [41] and VarTrix [42], should also provide an acceptable initialization.

#### 4.1.6 Distributions and Objective

Having introduced our model variables, we now summarize their respective probability distributions in Table 1.

Following this, we define our objective function, *L*, for the Expectation-Maximization (EM) algorithm. The reader is encouraged to refer to the table of definitions in conjunction with 2B. We note that we work in log-space for numerical stability and ease of computation.

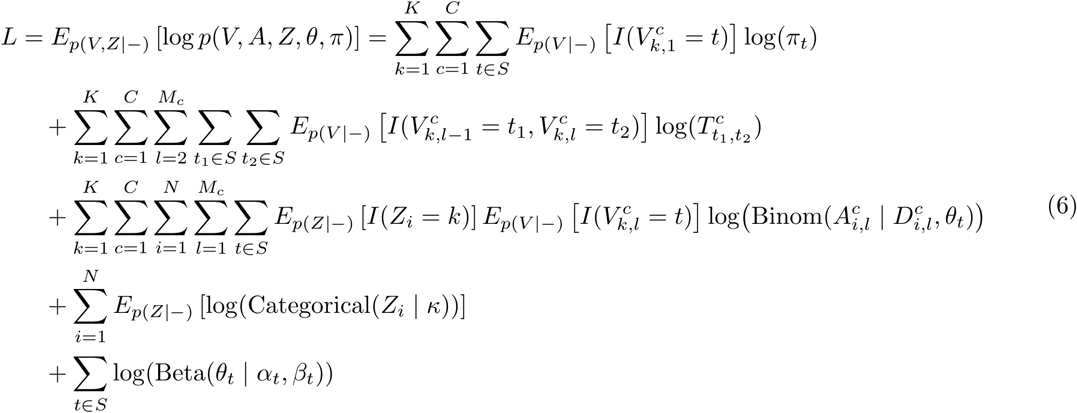

From this objective, we define the following quantities, which we will later have to evaluate:

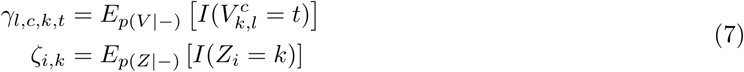

We can then rewrite the objective using these definitions to obtain the form we will work with for the update steps.

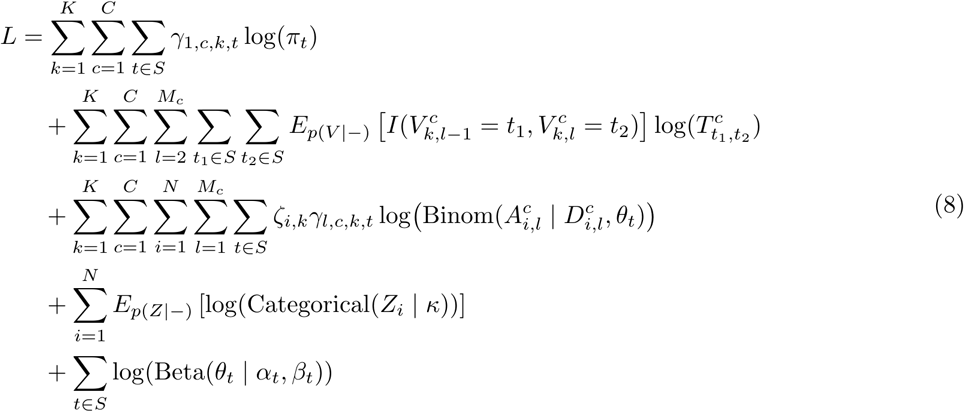

By differentiating the objective function with respect to each variable and its constraints, we can find each update step. We now present the workings for each of these updates.

#### 4.1.7 *Z* update

If we exponentiate our objective function we notice that:

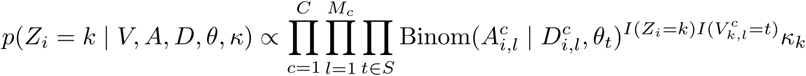

Then, we define:

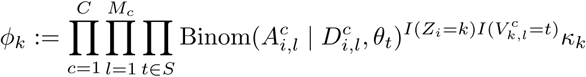

Normalizing gives:

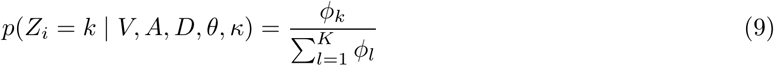

In fact, we can compute some of our previously defined quantities:

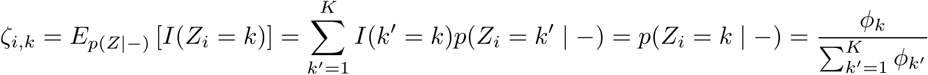

As we may want to work in log-space with *?*_*i,k*_, we quickly see that

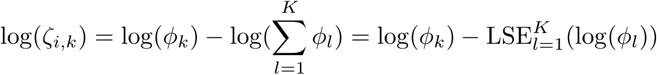

where 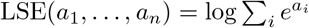.

#### 4.1.8 *π* Update

We consider the objective terms that depend on *π* only and differentiate, while also enforcing that *π* is a probability vector.

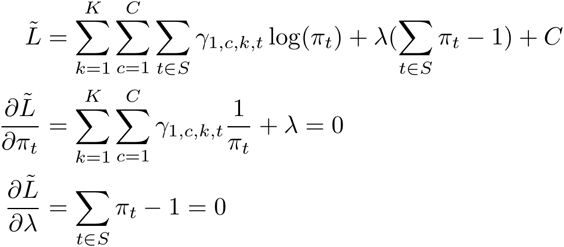

Solving for *π*_*t*_ and *λ*, we get:

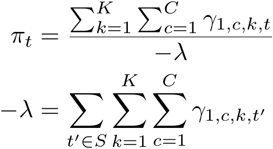

The above gives us an update of:

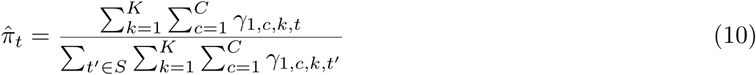

#### 4.1.9 *θ* Update

We restrict the loss function to terms that depend only on *θ*.

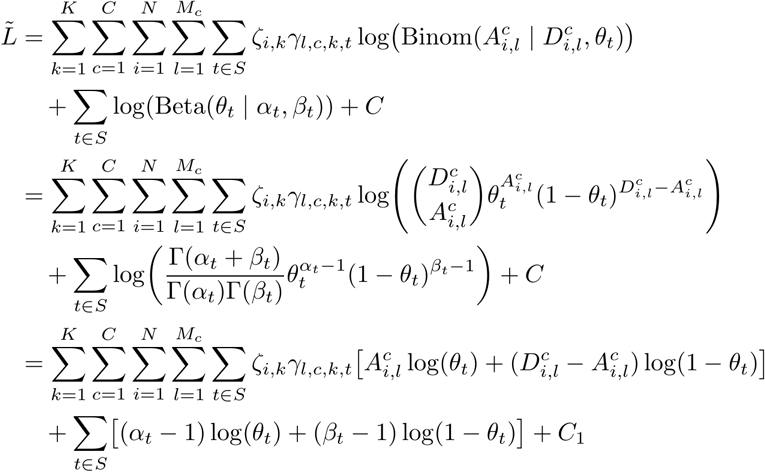

Setting the derivative to zero gives:

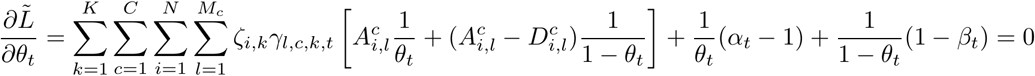

Rearranging, we get:

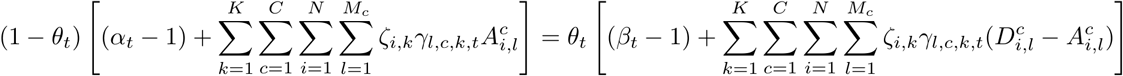

This gives us an update step of:

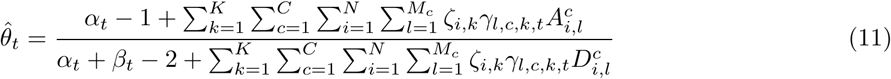

#### 4.1.10 *V* update

We now focus on inferring the values of *V* (our chains) through a forward-backward algorithm. Since 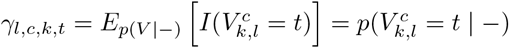, inferring *V* will also let us compute *γ*. First, we let *I*_*k*_ be the set of indices of cells assigned to cluster *k*, i.e. *I*_*k*_:= {*i*: *Z*_*i*_ = *k}*. Then, we observe that:

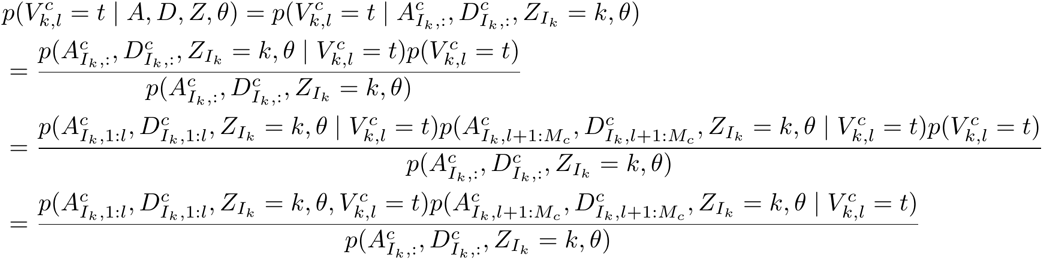

We then define

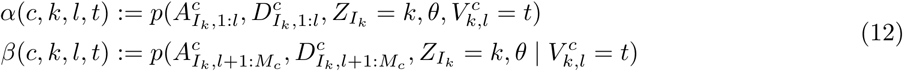

We then move our objective to log-space for stability.

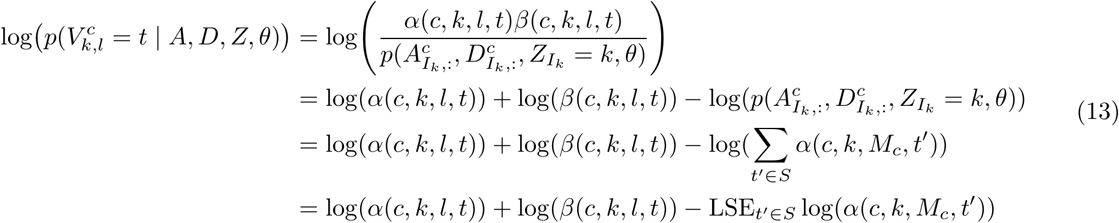

We can compute *α* and *β* recursively. For *α*, the computation is as follows:

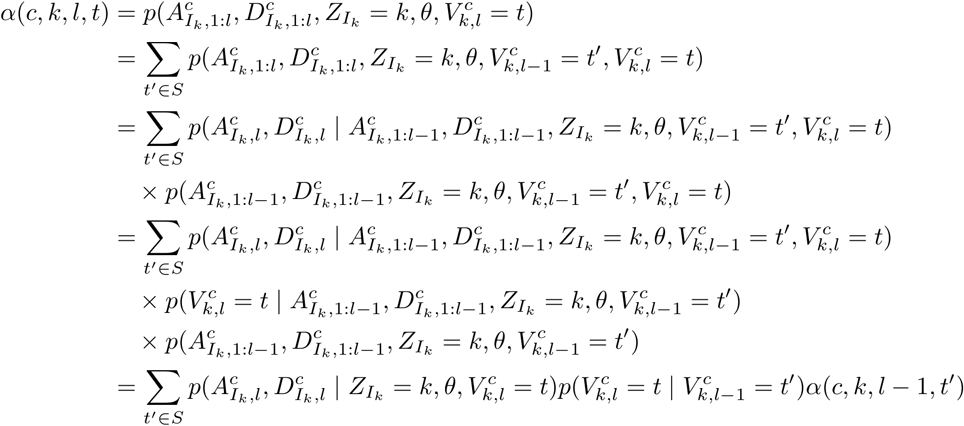

In log-space this is:

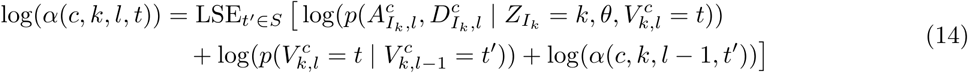

We initialize *α* as:

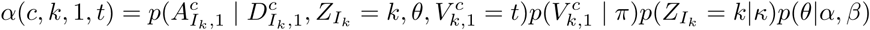

We can find *β* in a similar recursive fashion as follows:

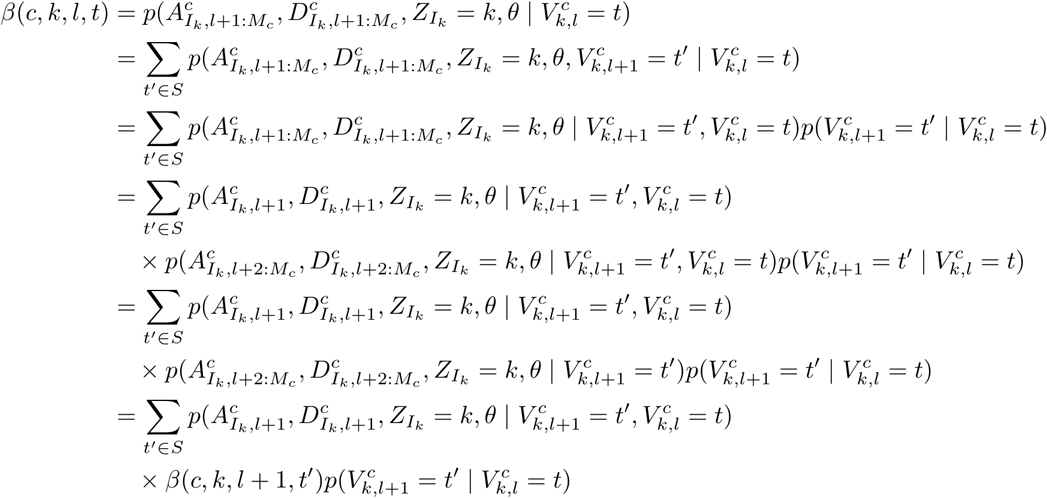

In log-space this is:

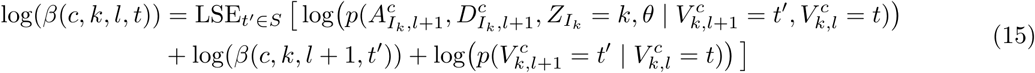

where we initialize *β*(*c, k, M*_*c*_, *t*) = 1. With these recursive formulas, we can then compute the maximum a posteriori values of *V*. Additionally, plugging in our values of *α*(*·*) and *β*(*·*) into Eqn. 13 allows us to compute values of *γ*_*l,c,k,t*_ as needed for the other updates.

### 4.2 Convergence Criterion

We now present a computationally cheap method for determining convergence. Our goal is to compute an objective function at each iteration with minimal extra work, determining convergence when its value changes minimally. The obvious choice would be to use our defined objective function, *L*. However, in practice, it is computationally expensive to calculate it at each iteration. Instead, we settled for the log joint probability of the current genotype sequence, which we denoted as *J* for simplicity. If we denote the log joint at iteration *i* as *J*_*i*_, we can take the difference at iteration *i* as Δ_*i*_:= |*J*_*i*_ *− J*_*i−*1_|. Convergence is then said to be achieved when Δ_*i*_ *≤ δ*, where *δ* is some user-defined threshold. Unlike *L, J* can be recovered using previously computed variables. We recall that we defined 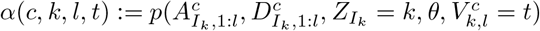 (with dependence on *π* encoded in the definition of *α*(*c, k*, 1, *t*)). This definition yields a natural way to compute the joint of the current sequence as:

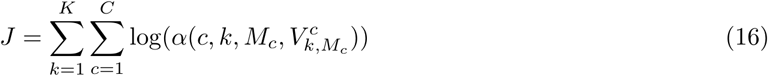

Since *α* is computed for each update, this provides an effective way to compute an objective function for convergence.

### 4.3 Consensus Clustering

It is common for EM methods to perform multiple restarts with differing initializations, only keeping the one with the best model probability. In the case of DemuxHMM, one might notice that the “worse” restarts are often only slightly worse in terms of ARI or homogeneity. We hypothesize that the worse restarts may still contain valuable clustering information. Then, on the whole, we might imagine that taking an average (or *consensus*) clustering may yield a stronger clustering overall. This idea of consensus clustering is a crucial post-processing step in the DemuxHMM model to avoid wasting EM restarts and to boost performance for low cost.

We implemented an efficient consensus clustering algorithm that can scale to thousands of clusters. For reference, Ghosh and Acharya provide a good summary of existing approaches in [43]. For our use case, we found a cumulative vote approach to be simple and effective. That is, we aligned cluster labels from each restart to a reference clustering (the best restart by model probability). Then, the mode label for each cell is its consensus cluster.

Mathematically, let the reference clustering be *R*, where *R*_*i*_ is a cluster label for cell *i*. Then, let *C*^*r*^ be a clustering for repeat *r* where 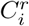 is another cluster label for cell *i*. We computed a Jaccard similarity between each pair of labels in the reference and repeat. The Jaccard similarity between labels *j, k* is defined as:

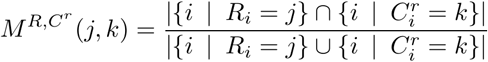

Then using 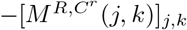 as the cost in the Hungarian algorithm [44], we can create the optimal one-to-one label matching to the reference. Iterating this approach for each *C*^*r*^, we constructed a cells-by-clusters matrix, *H*, where *H*_*i,r*_ is the reference-aligned-label for cell *i* in clustering repeat *r*. Taking the mode label for each cell then yields a consensus clustering.

In practice, we found the consensus clustering step to be fast to compute, taking a small fraction of overall compute time. We observed that it yielded noticeably improved performance in almost all test datasets.

### 4.4 Doublet Detection

Doublet detection is a natural extension to our SNP-based demultiplexing algorithm. Here, we show how DemuxHMM can be extended to compute the probability that a cell is a doublet. Let the variable *d*_*i*_ *∈* {0, 1} be an indicator for whether cell *i* is a doublet. Using Bayes’ rule we can create a workable form for *p*(*d*_*i*_ = 1|*A*_*i*_, *D*_*i*_), where *A*_*i*_ and *D*_*i*_ are the observed SNP data for the cell.

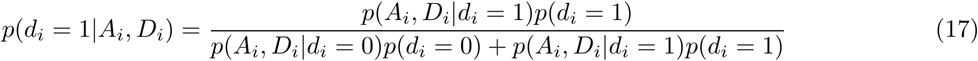

Here, *p*(*d*_*i*_ = 1) is a prior on the doublet rate of the data. Doublet rates tend to depend on the concentration of cells sequenced [37, 45]. However, an uninformative prior can set *p*(*d*_*i*_ = 1) = 0.5. Naturally, *p*(*d*_*i*_ = 0) = 1*− p*(*d*_*i*_ = 1). Additionally, we can take the data probability given a singlet to be our pre-existing cluster probability. That is, suppose that the model predicts the best cluster for cell *i* to be *k*, then:

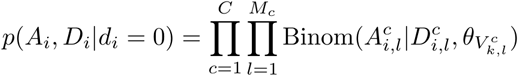

The above quantity is already computed as part of the inference steps. Thus, the doublet detection challenge stems from defining *p*(*A*_*i*_, *D*_*i*_| *d*_*i*_ = 1).

To compute the data likelihood given a doublet state, we make the assumption that the emission frequency of a doublet allele is the average frequency of its constituent singlet clusters. More precisely, suppose cell *i* is a doublet from clusters *k*_1_ and *k*_2_. Then, the emission frequency of the alternate allele of SNP *l* on chromosome *c* is:

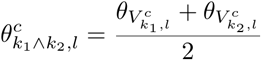

This emission probability can then be substituted into our binomial emissions formula. Of course, there may be multiple candidate clusters for the doublet source. We considered the top *q* most likely clusters, and weighed each possible pairing equally. These clusters can be chosen based on the best singlet cluster probabilities. Then, using our binomial emission probabilities, we can say that:

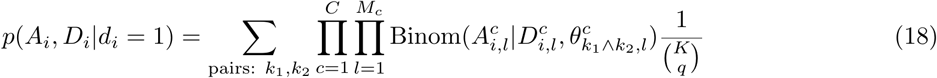

We note that in practice, it is helpful to compute these binomial products in log-space.

Now, we have the machinery to evaluate Eqn. 17 on a cell-by-cell basis. Eqn. 17 gives us the ability to predict the probability that a cell is a doublet. Doublet cells should be filtered out before downstream applications. Additionally, after filtering, users can then refine the demultiplexing results by performing a second doublet-free pass. We evaluate and discuss doublet detection performance in Sec. 2.3.

### 4.5 Pre-processing for the Diversity Outbred Mice Dataset

In this section, we describe the pre-processing steps used to prepare the pooled Diversity Outbred (DO) mice dataset for demultiplexing from Dillard et al. [22]. The dataset comprises scRNA-seq data from the bone marrow of 77 mice bred using the diversity outbreeding scheme. DO uses eight founders in a breeding scheme that leads to contiguous SNP blocks compatible with our model. The Dillard et al. dataset pools mice into batches of four or five, leading to a partial ground truth. We preserved these pool identities for later use. However, we processed the dataset as if all mice were pooled together.

We now describe the mapping, variant calling, and filtering steps. To start, we mapped the reads to the GRCm39 reference genome using cellranger 7.2.0 [37] with default settings for quality control. After the mapping, 116,890 cells remained. Next, we merged all pools of mice. We used cellsnp-lite 1.2.3 [40] with parameters `--minMAF 0.1 --minCOUNT 20` to call variants on each chromosome. Rather than calling all variants de novo, we called against the collection of mouse SNPs associated with [46]. This still resulted in over 2.3*·*10^6^ sites with non-zero calls, necessitating filtering to be usable. To start, we filtered variants to only those reported on at least 20 cells (depth *>* 0). Next, we computed a Fisher information score based on alternate allele fraction. Consider variant *l* on chromosome *c*. We can take the alternate allele fraction as:

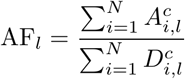

As variant draws are 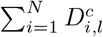 Bernoulli trials, the Fisher information is given as:

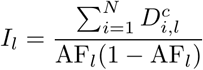

We kept the top 1% of SNPs on each chromosome using *I*_*l*_, leaving a total of 23,252 variants. Next, we recognized that the filtering may have impacted the usability of some cells. To adjust, we filtered out cells that had a variant depth of less than 100 UMI and those that did not have an alternate allele UMI on each chromosome. The cell filtering left 86,971 cells and was the final step in our pre-processing pipeline. Thus, our demultiplexing-ready dataset had 86,971 cells and 23,252 variant sites.

For additional testing, we created read-downsampled datasets for mouse. To keep variants consistent between downsamplings, we downsampled the variant count matrix, *A*, and the reference count matrix *R* = *D − A* directly. To do this, we re-sampled every non-zero entry as Binom(*·, f*), where *f* is the fraction of reads we wanted to keep. We re-sampled for *f ∈ {*0.1, 0.25, 0.5*}*.

### 4.6 Demultiplexing on Diversity Outbred Mice

To start, we needed to create a good estimate for the transition matrices *T*^*c*^. For estimating *T*^*c*^, we first needed an estimate of the expected number of state transitions in DO mice by generation. Svenson et al. measure a median of 617 recombination points for generation 20, with a predicted increase of 28 per generation [27]. Based on data provided by the authors of [22] (see Supplementary Table 1), we computed that the mice used had a mean generation of 25.86. Extrapolating the number of recombination points as *b*(*g*) = 617 + 28(*g −*20), we got *b*(25.86) = 781. Assuming these points were distributed roughly evenly by chromosome pair, we got 39.05 expected transitions per pair. This number then allowed us to estimate *T*^*c*^ as described in Sec. 2. We used an *ϵ* = 0.01 to account for errors in estimation.

Next, we set initial values for *π*, and *θ*. To do this, we ran DemuxHMM with 60 re-initializations and chose the one with the best model probability to set the parameters. This model gave us a *π* = [0.735, 0.192, 0.073], and *θ* = [0.015, 0.445, 0.963].

Finally, we demultiplexed using these parameters. We performed two-pass demultiplexing with 240 re-initializations on each pass. The first pass fit the model on the full dataset and detected doublets. We considered cells doublets if the model marked their doublet probability as above 90%. We assumed a doublet prior of 5% (also provided to all competing methods). The second pass worked on the dataset after we filtered the detected doublets out. The clustering was initialized as the consensus clustering of the doublet-containing runs (with doublet assignments removed). We found that a single EM restart was sufficient for the second pass, as most restarts converged to the same solution.

### 4.7 Heuristic parameter selection for K

A natural question that arises when fitting the model is how to choose the number of individuals, *K*. For the examples we studied in the results section, we had a ground truth available to us. When this ground truth is not available, we suggested a heuristic method based on the objective *J* given in Sec. 4.2. The heuristic approach relies on plotting *J* as the number of clusters varies in a reasonable range. We found that as the number of clusters reached, and surpassed the true *K*, the curve of corresponding *J* values flattened near some limit. This effect can be seen in Fig. 9A on the DO mouse dataset with a true *K* = 77. Similarly, this effect is also clear in Fig. 9B on a set of 6 resamplings of simulated Drosophila data with a true *K* = 50. A user should then select a value for *K* in the flattened area of the curve. In our experience, the model behaves well if *K* is over-estimated, incurring no significant clustering performance penalty. The main penalty we noted for over-estimation is an increase in runtime. Thus, this technique serves as a simple heuristic to select *K* when it is unknown.

**Fig 9.**
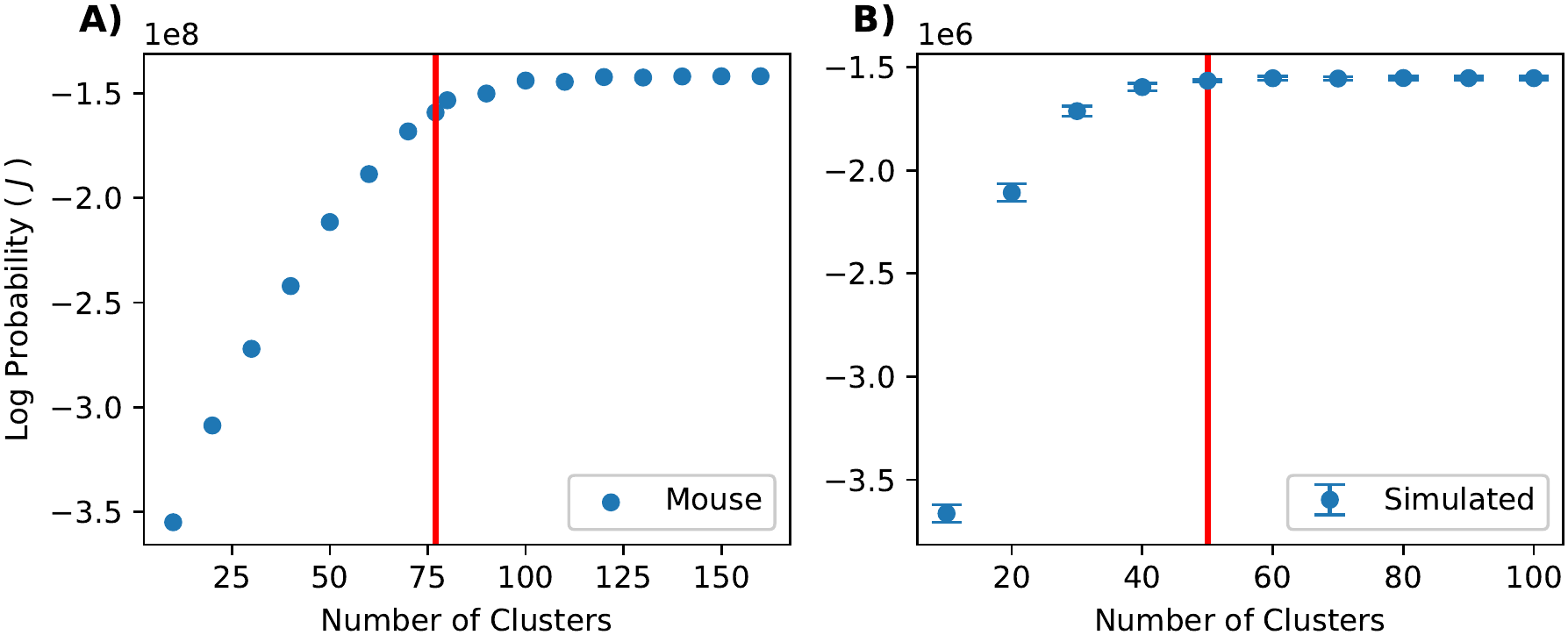
A plot of the model objective, *J*, as the number of clusters varies. The objective is shown for the real DO mouse dataset (A), and a simulated Drosophila dataset (B). In panel A, points are the model probability from the best EM restart (of 30) on the full DO mouse dataset. The red line indicates the true number of clusters at *K* = 77. In panel B, points are a mean of 6 repeats over resamplings (again taken as the best EM restart of 30) of the Drosophila datasets used for Fig. 5 with a true *K* = 50, indicated by the red line. Error bars indicate standard deviation.

**Fig 10.**
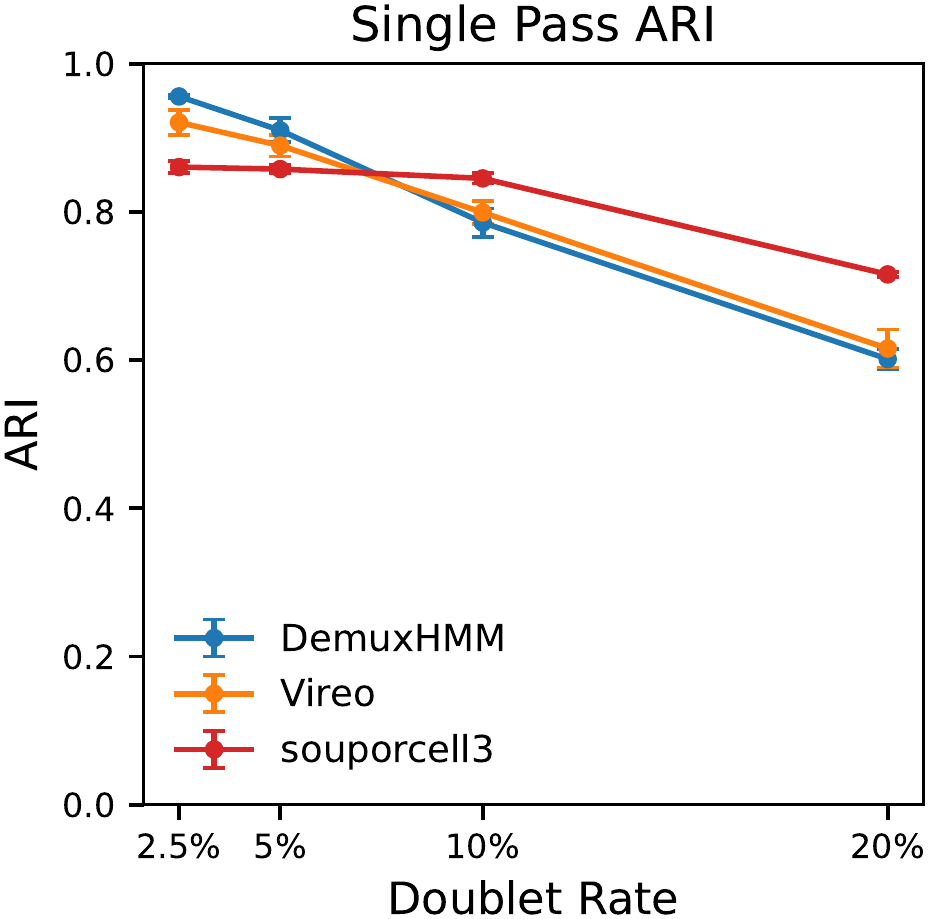
A comparison of method performance on 50-individual, 20,000-cell (pre-quality-control) *Drosophila* datasets with varying doublet rates. ARI is given without doublet detection or any removal of doublets. Values are a mean of 6 data resamplings. Error bars indicate standard deviation. scSplit is not included as it does not assign clusters to doublet cells, which skews its performance downward.

**Fig 11.**
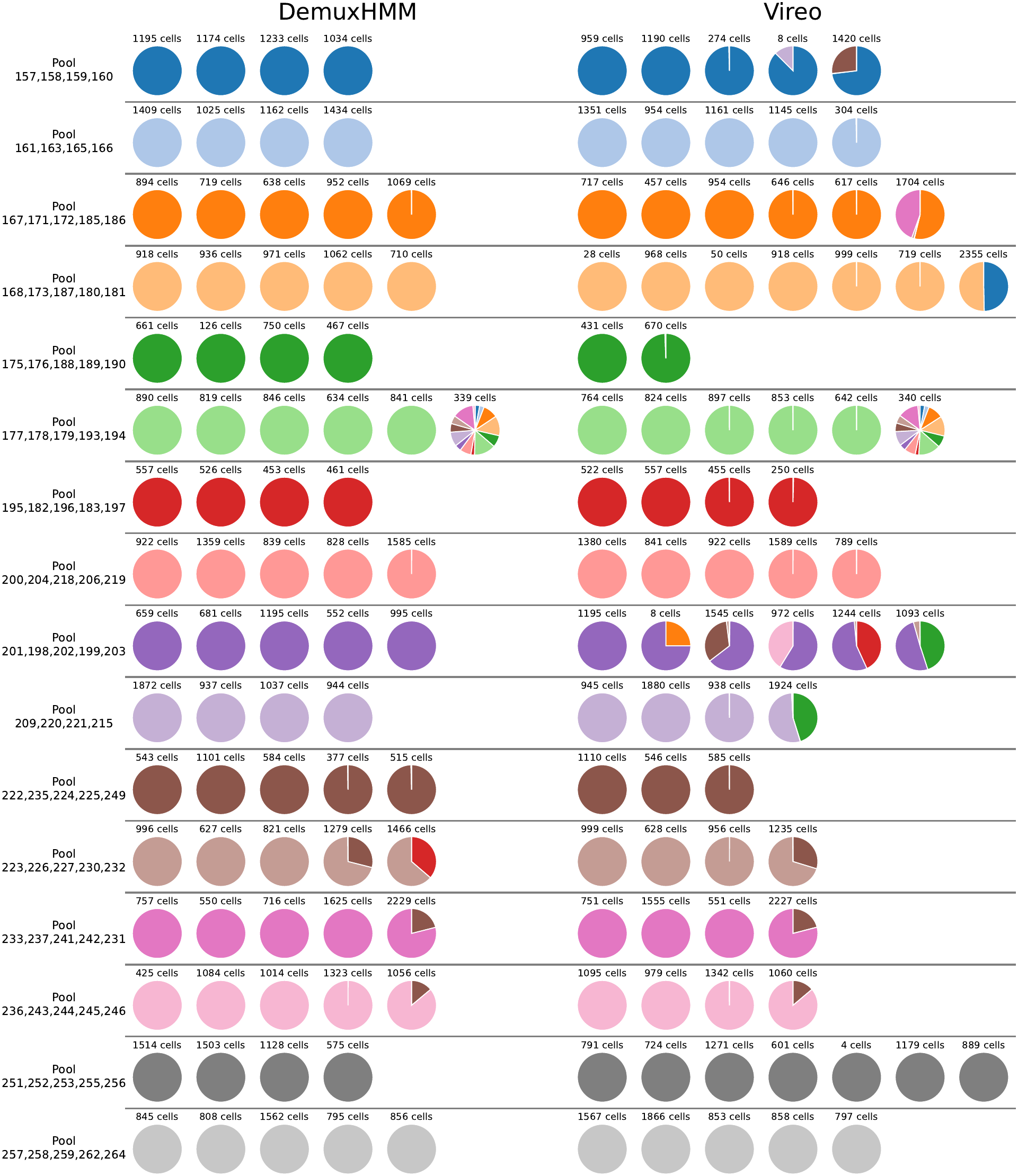
A view of clustering on the DO mouse dataset for DemuxHMM compared to Vireo. Each cluster is represented as a pi chart and annotated above by the number of cells contained within. The colors of the pi chart represent the cluster’s composition with regard to ground truth pool identity. We assign clusters a row based on majority pool fraction. Pool names represent the lab-assigned numbers for mice sequenced in that pool. Clusters for both DemuxHMM and Vireo shown are the result of two-pass demultiplexing, which includes doublet removal.

### 4.8 Synthetic Data Generation

We generated a synthetic dataset that simulates our suggested experimental structure from Section 1. Our starting data is a list of DSPR A4 strain single nucleotide polymorphisms (SNPs) relative to DSPR A6. We filtered this list to SNPs on the top 15% of genes by variant count. We found that genes with more than one variant gave stronger signal for demultiplexing. This list was then filtered to SNPs that appeared in the last 150 base pairs of a gene. These are the SNPs that can potentially be captured by common scRNA-seq technologies.

Next, we simulated a pool of individuals. An individual is seen as a set of chromosomes and locations of SNPs along them. We assumed that the F0 generation contained a pure A4 male, and a pure A6 female. For each generation, we performed the following steps until the desired number of offspring was simulated:

1. Choose one male and one female breeding pair for the pool.
2. Simulate meotic recombination in the chromosomes. The number of recombinations and their locations are determined by the recombination rate calculator [32]. Chromosome 4 is assumed to be conserved.
3. For each chromosome, choose which chromatid goes to the offspring from each parent.
4. Randomly assign the gender of the offspring.

We repeated this procedure until the desired number of offspring had been simulated to form the next breeding pool. This was done for *G* generations.

Next, we simulated the resulting variant call matrices. We assumed that we saw *N* cells from each individual. We sought to generate matrices *A*^*c*^ and *D*^*c*^ that have cells on rows and variants from chromosome *c* on the columns. Entries in *A*^*c*^ represent alternative allele calls for that cell and variant. Entries in *D*^*c*^ are the overall depth for that cell and variant site. For each of these cells, we performed the following procedure.

1. Sample a scRNA-seq cell profile, 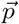, from the combined datasets of the head and body from the Fly Cell Atlas [33]. We chose only cells that had a minimum of 1,500 UMI counts to ensure our sample was of high quality.
2. We normalized this profile 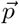, to the probability vector 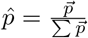.
3. We sampled *U* UMI counts, using 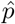 as the probability of drawing a count from a gene.
4. For each count, randomly select which of two homologous chromosomes it came from. If the chromosome has a detectable variant on that gene, increment the number of alternative allele calls for that cell/variant combination by one, and also increment the depth at that variant. If the chromosome is *X*, then males will have only one chromosome to sample from.

Following this procedure, we obtained matrices *A*^*c*^ and *D*^*c*^, which we could use to evaluate the methods. To maximize performance across all methods, we also filtered these matrices to only cells that had variant reads on all three non-conserved chromosomes (X, 2, 3). Generally, variants on the X chromosome are the limiting factor as our test dataset contained approximately 25% as many variant sites compared to chromosomes 2 and 3. In practice, this filtering step kept approximately 50% of cells or more, depending on the number of breeding generations used. We do note that all methods produce reasonable results without filtering, though with reduced performance. Thus, in real datasets, groups may choose to forego this step if they deem the performance trade-off worth the additional cells.

#### 4.8.1 Simulating Doublets

To create a dataset with simulated doublets, we modified the procedure from above. As a doublet is the combination of two cells, we can assume that its allele counts are also combined. Supposing that we want *N* total cells with a doublet rate *r*, we need to create (1*− r*)*N* singlets and 2*rN* additional cells that will be merged into doublets. Cells to be merged are paired uniformly at random. Suppose cells *i* and *k* are to be merged into a doublet. The doublet cell has alternate allele counts 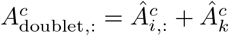 and depths 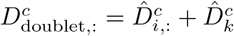, where *Â*^*c*^ and 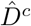 indicate pre-merger matrices. To create Fig. 8, we simulated 50 individuals with 20,000 total pre-filtering cells across all individuals. Doublet rates of 2.5%, 5%, 10%, and 20% were used. 6 resamplings of the datasets were created for bootstrapping error.

### 4.9 Demultiplexing on Drosophila Datasets

Here we detail the settings used to demultiplex on Drosophila. The majority of Drosophila tests used the same settings and parameters. Importantly, *T*^*c*^ was computed using a large, representative set of 1,000 individuals at the relevant number of breeding generations (22 for all but Fig. 7A). As such, we used a small *ϵ* = 10^*−*9^. The number of individuals, *K*, was set to the number of pre-filtering individuals for all methods. The logic behind this choice was that this is the number that would be available to a researcher demultiplexing a new dataset. *θ* was taken to be [0.006, 0.469, 0.988], computed in an initial training run. We did not specify a *π* initialization and instead learned it on the fly. With regard to EM repeats, 30 were found to be sufficient for all DemuxHMM tests, except the high-individual-count experiment, in which 60 were used. However, we do note that there is room to improve results with more repeats due to the consensus clustering approach. As the Drosophila datasets are smaller than mouse, we were able to run multiple restarts on the GPU concurrently. 20 processes were used when below 500 individuals, and 10 otherwise. Finally, for the doublet-containing tests in Fig. 8, the doublet prior provided (to all methods) was the ground truth fraction of doublets.

### 4.10 Impact of Demultiplexing Error on Trajectory Inference

We simulated the effect of demultiplexing error on trajectory inference by shuffling time labels in a *Lytechinus variegatus* (sea urchin) dataset [36]. Specifically, the dataset has 18 timepoints (collections of urchin embryos) in the range of 2-24 hours post-fertilization (hpf). We chose ratios of cells *x*∈ [0, 0.9], such that for each such *x* we took *x*% of cells at each timepoint and randomly reassigned their time label. We quantified ARI for each level by measuring against the original labels. These datasets were meant to represent demultiplexed datasets with a full spectrum of possible ARI errors. For each dataset, we performed trajectory inference as described in the original study. The output of this inference was a set of cell fate matrices for each shuffling ratio and timepoint *t*, which we denoted as *F*^*x,t*^. If the shuffled dataset has *N*_*x,t*_ cells at time *t*, then *F*^*x,t*^ is an *N*_*x,t*_*×* 6 matrix. A row, 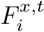, is a probability vector indicating the likelihood that the cell becomes one of the 6 main cell types at the terminal time of 24 hpf. To quantify error, we focused on the difference of *F*^*x,t*^ from baseline, *F*^0,*t*^, for *t* = 10. To do this, we used the Earth Mover’s Distance (EMD) [47] with a cost matrix of *L*^1^ norms between rows. That is, the cost matrix for shuffling level *x* is given by 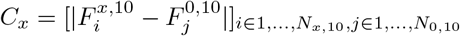. Each row is weighted uniformly. The resulting EMD error can be roughly interpreted as the amount of probability that has shifted between entries in each row. We also qualitatively analyzed changes in fate probabilities by projecting the probability of a cell to become the SMC cell-type or endoderm, versus any other cell type in a 2D barycentric representation. These errors and barycentric projections are illustrated in Fig. 6.

### 4.11 Competing Methods Demultiplexing Settings

#### 4.11.1 Souporcell3

Souporcell3 (release version 3.0) was run using 60 threads. Souporcell3 features were turned on (`-s true`), and the `-m khm` flag was used (as recommended for large datasets). Datasets were written using a custom function which included some zeros in the matrix market files to preserve alignment. This differs from the typical sparse representation, but does not alter the stored data. Souporcell3 appears to incorrectly read files where these zeros are not explicitly saved, resulting in poor performance.

#### 4.11.2 scSplit

scSplit version 1.0.9 was run with default settings. Minor modifications were made to the codebase to support updated versions of Python libraries. Additionally, a small `eps=1e-10` term was added to the cluster probability calculations to avoid divisions by zero. We do not expect these changes to impact the fundamentals of the approach.

scSplit occasionally does not assign cells if they don’t meet an internal probability threshold. In doublet-free testing, we treated these as incorrectly clustered. However, our experience shows this only affects a maximum of a few cells and has no practical impact on performance.

#### 4.11.3 Vireo

Vireo version 0.5.9 was run with default settings except for increases to the number of maximum iterations. We set these to `max_iter=5000, max_iter_pre=2500` to tackle occasional slow-converging cases. 50 initializations were used, which is the default. However, we do note that the parallelization for these was broken in the current version of the package, so we re-implemented it as faithfully as possible. The maximum 50 threads were used for Drosophila, while memory constraints limited us to 10 threads for mouse.

## 5 Supporting information

### 5.1 S1 Appendix - Related Work

We divide existing demultiplexing techniques into those that still require some manual manipulation of individuals and those that do not. For the application of creating a massively high-resolution time course, as we need to eliminate individual handling bottlenecks, the latter techniques are most relevant. Nonetheless, they still have performance limitations when there are many individuals or when there are few cells per individual.

Roughly, there are two categories of methods: (1) methods that experimentally tag individuals, (2) computational methods based on natural genetic variation “barcodes”. Experimental methods generally require individual-level handling for an initial molecular barcoding step. These methods do not require any inference, and hence, errors stem mainly from tagging quality. This can be useful for collecting datasets where variant sites may be limited, or sequencing depth is insufficient at those sites. For example, studies involving rare cell types may benefit from this approach. Computational methods, in contrast, typically perform a probabilistic inference based on profiles of single nucleotide polymorphisms (SNPs) that are unique to each individual. The probabilistic nature of these approaches leaves them open to the possibility of assigning some cells incorrectly. We further subdivide the computational approaches into those that require prior information on the genotypes of the individuals and those that do not. Genotype-requiring techniques have high accuracy as they do not have to infer each individual’s genotype. However, they do require individual-level bulk sequencing or similar, leading to the same bottleneck. Thus, when parallelization is key, we must turn to a class of methods we will refer to as self-genotyping. Self-genotyping methods allow researcher to sequence all individuals in a single pool with no individual-level processing. These methods only require a reference genome for the species strain as a whole to map reads and call variants, but they do not need genotypes of individuals for demultiplexing. However, they carry the added challenge of needing to infer both genotype and cell identity, leading to lower quality assignment when conditions are not optimal.

#### 5.1.1 Experimental Tagging of Individuals Using Molecular Barcodes

We first discuss experimental techniques which tag individuals with molecular barcodes. The most common involve either some form of cell hashing or a split-pool style combinatorial barcoding approach. There are a number of existing cell hashing methods. The overall approach involves performing individual-by-individual reactions to introduce a barcode for each individual. The approaches to achieve this are varied. TotalSeq uses HTO Antibodies to deliver barcodes [8]. On the other hand, MULTI-seq [10], 10x CellPlex [11], ClickTags [12, 13], and SUM-seq [9] use modified oligos to deliver barcodes. These methods generally require a technician to perform reactions on each individual, substantially limiting scalability. Split-pool techniques aim to improve this bottleneck with combinatorial indexing. One loads individuals into wells which receive unique barcodes. Then, one pools them and repeats the process multiple times. Each pooling and barcoding step expands the potential barcode space exponentially. The first barcoding step applies barcodes at the individual level. Two prominent split-pool methods are sci-rna-seq3 [48] and SPLiT-seq [14]. Both techniques still require some individual-by-individual processing, at minimum, pipetting each into a separate well and adding reagents well-by-well. With larger datasets, one may even need to split individuals among multiple wells, compounding this effort. Split-pool techniques generally offer better scalability than cell hashing, and have resolved large datasets such as Qiu et al.’s scRNA-seq mouse dataset containing 83 embryos [49].

However, due to the need to perform some individual-level work, split-pool methods only serve as a partial solution to creating high-resolution time courses, making it difficult to create datasets with hundreds or thousands of resolved individuals.

#### 5.1.2 Biological Tagging of Individuals Using Combinations of Natural Variants

We now explore a class of methods that use combinations of variants as barcodes. These methods assume that researchers pool individuals from the start and sequence them together, requiring no per-individual molecular tagging. Any single-cell sequencing technology which allows the calling of variants suffices. Single-cell pileups of variants serve as the input to these methods. Specifically, they seek to differentiate individuals by their profiles of SNPs. Since most naturally occurring SNPs are inherited and present in all cells of an individual, cells within the same individual share a similar SNP profile. If there is sufficient genetic diversity among individuals and high enough sequencing depth to capture it, then these SNPs form a lossy barcode that one can use to cluster cells back into individuals. We can split the class of methods into two categories: genotype-requiring methods, and self-genotyping methods. The two categories differ based on whether they require genotypes for each individual they demultiplex, with genotype-requiring methods requiring individual-level handling to obtain these genotypes.

Genotype-requiring methods offer a straightforward approach to demultiplexing with variants, at the cost of labor to obtain initial genotypes. These methods assume that the researcher knows the genotype of each individual in the pool a priori alongside single-cell variant pileups. The methods then focus on performing inference on a cell-by-cell basis to match the cell’s SNP profiles to one of the references. One can obtain genotypes through SNP microarrays [50], bulk sequencing, or other technologies. The three main methods in the genotype-requiring category are Demuxlet [15], Dropulation [16] and Demuxafy [17]. While having reference genotypes can improve accuracy, the need for individual-level processing can make these methods infeasible for datasets with individual counts in the dozens or greater.

Self-genotyping methods attempt to address this issue by inferring genotypes of individuals probabilistically. They can work purely from variant pileups of pooled cells. Generally, they approach the inference problem by probabilistically inferring genotypes from initial cell assignments, re-assigning cells based on those genotypes, and repeating until convergence. In this category, there are four main methods: Vireo [18], Souporcell3 [19], scSplit [20], and Freemuxlet [21]. The main advantage of these methods is that no individual-by-individual pre-processing is necessary. Removing this constraint theoretically allows for very high scalability. However, in practice, the inference of genotypes combined with demultiplexing is challenging. Various factors such as sequencing depth, genetic diversity, and SNP frequency can limit performance. Most notably, this results in a trade-off between the number of individuals we can demultiplex and the number of cells per individual.

## Acknowledgments

We thank Kathleen Griffin for her help with the design and preparation of Fig. 1 and Fig. 2A.

